# Placental-brain axis in females detected within broadly impacted metabolic gene networks protects against prenatal PCB exposure

**DOI:** 10.1101/2024.07.14.603326

**Authors:** Kelly Chau, Kari Neier, Anthony E. Valenzuela, Rebecca J. Schmidt, Blythe Durbin-Johnson, Pamela J. Lein, Ian Korf, Janine M. LaSalle

## Abstract

**Background:** Neurodevelopmental disorders have a strong male bias that is poorly understood. Placenta is a rich source of molecular information about environmental interactions with genetics (including biological sex), that affect the developing brain. We investigated placental-brain transcriptional responses in an established mouse model of prenatal exposure to a human-relevant mixture of polychlorinated biphenyls (PCBs).

**Results:** To understand sex, tissue, and dosage effects in embryonic (E18) brain and placenta by RNAseq, we used weighted gene correlation network analysis (WGCNA) to create correlated gene networks that could be compared across sex or tissue. WGCNA revealed that expression within most correlated gene networks was significantly and strongly associated with PCB exposures, but frequently in opposite directions between male-female and placenta-brain comparisons. In both WGCNA and differentially expressed gene analyses, male brain showed more PCB-induced transcriptional changes than male placenta, but the reverse pattern was seen in females. Furthermore, non-monotonic dose responses to PCBs were observed in most gene networks but were most prominent in male brain. The transcriptomic effects of low dose PCB exposure were significantly reversed by dietary folic acid supplementation across both sexes, but these effects were strongest in female placenta. PCB-dysregulated and folic acid-reversed gene networks were commonly enriched in functions in metabolic pathways involved in energy usage and translation, with female-specific protective effects enriched in PPAR, thermogenesis, glycerolipids, and O-glycan biosynthesis, as opposed to toxicant responses in male brain.

**Conclusions:** The female protective effect in prenatal PCB exposures appears to be mediated by dose-dependent sex differences in transcriptional modulation of metabolism in placenta.

**Graphical Abstract:** 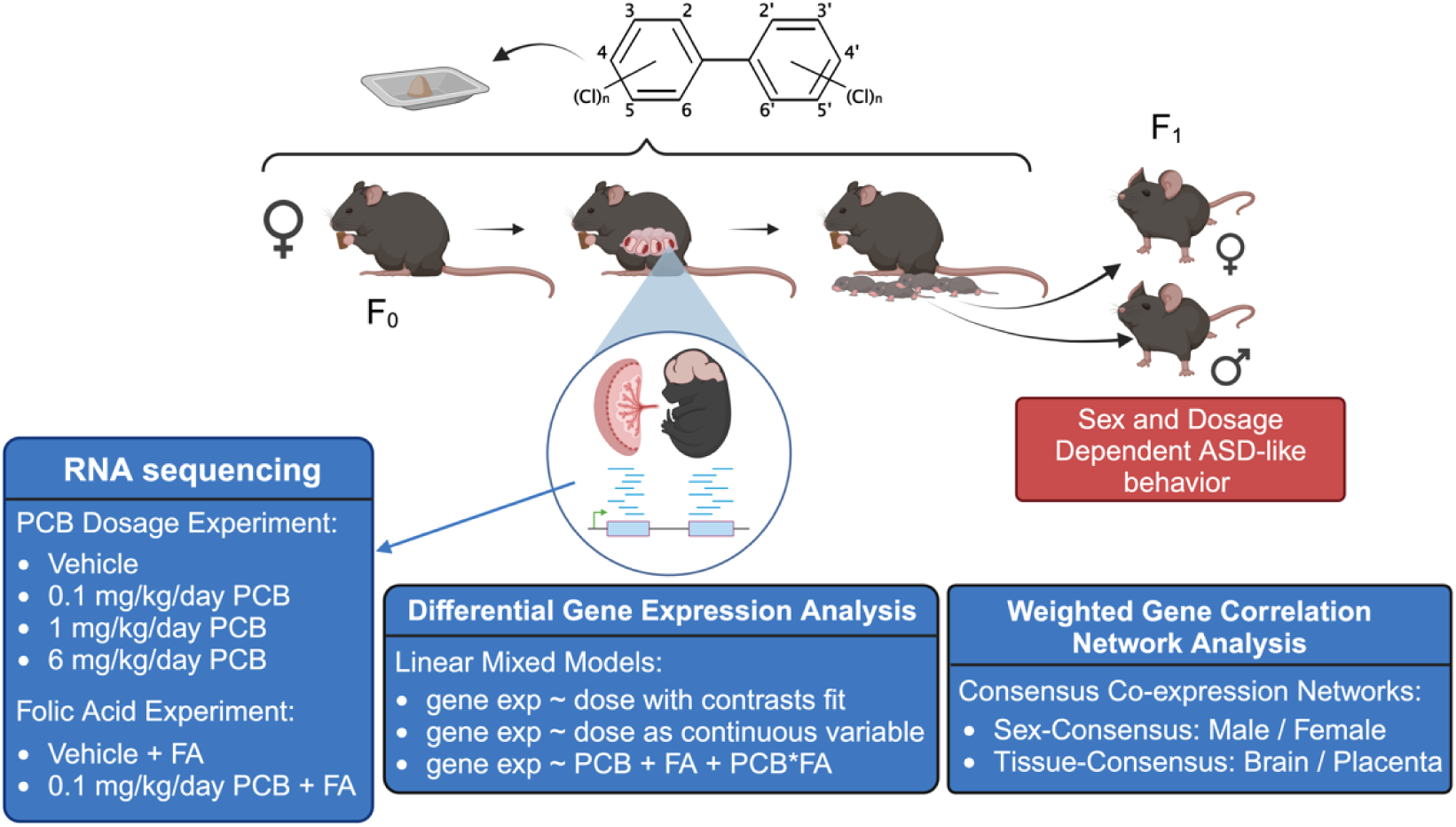

## Background

The placenta is a fetal-derived tissue that is critical for the development of eutherian mammals because of its interface with maternal nutrients and oxygen and serves as a barrier against external environmental factors. In humans, term placenta is accessible as a birth byproduct and potentially a rich source of signature gene expression biomarkers that can reflect the developing brain’s response to *in utero* exposures through a shared “placental-brain axis”. The placental-brain axis could be useful in investigating gene by environment interactions underlying the perplexing increase in the prevalence of neurodevelopmental disorders (NDD) of 1 in 5 males compared to 1 in 8 females in the US (2018-2021).^1^

Endocrine disrupting chemicals, including polychlorinated biphenyls (PCBs), are posited to be risk factors for male-biased autism spectrum disorders (ASD) and other NDD.^2, 3^ In the NDD enriched-risk human prospective study “Markers of Autism Risks in Babies - Learning Early Signs” (MARBLES), concentrations of PCBs detectable in maternal blood were associated with adverse neurodevelopmental outcomes including ASD and were associated with DNA methylation changes over known ASD risk genes.^4–6^ PCBs are present in the human brain, blood, and breast milk, and they can easily cross the placental barrier, exposing fetuses through maternal blood. Humans, including pregnant women and those of child-bearing age, encounter PCBs through inhalation and consumption of contaminated food and water. Exposure during pregnancy poses a significant risk for NDDs such as ASD.^7–9^ Interestingly, recent studies have found that folic acid supplementation during pregnancy is associated with reduced likelihood of ASD in the offspring, suggesting a protective effect against environmental exposures.^10–13^ It is therefore of interest to investigate folic acid’s protective effect in the presence of established neurodevelopmental toxicants such as PCBs.

In a mouse model of prenatal exposure to the mixture of PCB congeners detected in the MARBLES cohort, there were significant deficits in sociability and ultrasonic vocalizations, as well as enhanced repetitive behavior but only in males, specifically at the lowest dose tested (0.1 mg/kg/d).^14^ This observed male bias in the multitude of behavioral phenotypes prompts consideration of a “female protective effect” during PCB exposures experienced *in utero* that have been previously suggested in both genetic and environmental causes of NDDs.^15–17^ Furthermore, the non-monotonic dose response of behavioral phenotypes manifesting only at the lowest dose tested, rather than the vehicle, 1 mg/kg, or 6 mg/kg doses, is particularly compelling.^14^ A whole-genome DNA methylation analysis confirmed sex-specific differences in DNA methylation profiles and uncovered that the placenta and fetal brain share a NDD DNA methylation profile in a mouse model of prenatal PCB exposure.^18^ These findings prompted investigation of the complex interactions between the placental-brain axis, the observed sexual dimorphism, and the non-monotonic dosage effects.

To investigate possible sex differences in placental-brain transcriptomes, including effects of PCB dosage and folic acid supplementation, we exposed C57Bl/6J females to varying doses (0, 0.1, 1, 6 mg/kg) of a human-relevant PCB mixture representative of the PCB profiles in the human cohort MARBLES study. To test whether folic acid supplementation during pregnancy counteracts effects of prenatal exposure to PCBs, mice were supplemented with or without folate prior to timed mating. At embryonic day 18, brain and placenta were collected for RNA-sequencing. This mouse model was designed to identify tissue- and sex-specific gene expression signatures underlying the sexual dimorphism and non-monotonic dose response of the developmental neurotoxicity of PCBs, as well as the potential counteracting effects of folic acid supplementation during pregnancy.

## Results

### Experimental design to examine dosage effects of a human-relevant PCB mixture and potential protective effects of folic acid supplementation

Figure 1A shows the experimental design and experimental groups that were included to test the hypothesis that PCB-induced transcriptional changes in the placental-brain axis will be affected by both dosage and sex. Dams were dosed daily for 2 weeks before conception, through breeding and gestation, until embryonic day 18 (E18). The effect of dietary supplementation with folic acid was also investigated at a single low dose (0.1 mg/kg/d) of the PCB mixture. RNA was isolated from E18 matched placenta and brain samples for transcriptome analyses via RNAseq. We then used two complementary bioinformatic approaches to analyze RNAseq data. Traditional differential gene expression (DGE) analysis detects changes to individual genes in response to treatment and assumes that a small proportion of genes will change. In contrast, weighted gene correlation network analysis (WGCNA) is based on the biological principle that genes respond within correlated networks and pathways, thereby allowing pathway-focused rather than gene-focused analyses.^19^

**Figure 1.**
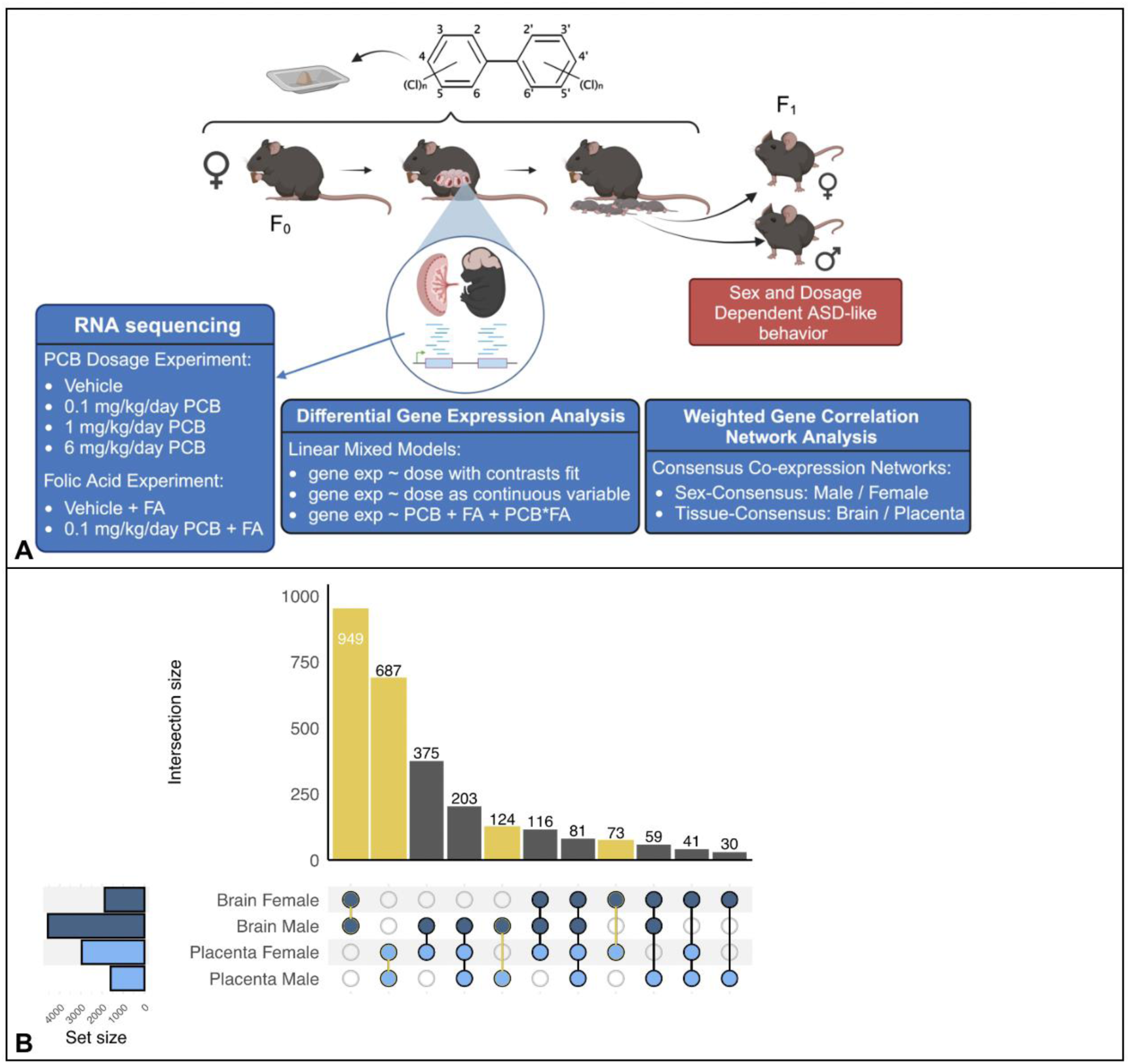
Experimental design and overlap in PCB dose-dependent differentially expressed genes (DEG) by sex and tissue. **A.** The MARBLES mixture of PCBs was given orally in daily doses preconception and prenatally until embryonic day 18 (E18) when placenta and fetal brain samples were harvested for RNAseq. There were two different experiments (PCB dosage and folic acid effect) and two different analysis approaches (DGE and WGCNA). **B**. Upset plot of DEGs (uncorrected p<0.05) identified based on PCB dose effects as a continuous variable shows that largest number of DEGs in male brain and the least overlap with female brain and other categories.

### Analysis of individual differentially expressed genes reveals complex sex, dosage, and tissue effects of PCB mixture and folic acid

We first used an analysis of individually differentially expressed genes using limma voom, a statistical pipeline designed for RNAseq data with multiple treatment groups. The results are summarized in Table 1 and Figure 1B which show strong effects of dosage, sex, and tissue type on the number of DEGs. For instance, the number of DEGs followed a primarily monotonic dose-response pattern in males, with the most differentially expressed genes detected at the 6 mg/kg/day dose level vs. vehicle control (318 genes) when PCB dose was modeled as a continuous variable (368 genes) (Table 1). In contrast, the largest number of DEGs from female brain was in the low dose (0.1 mg/kg/day) versus high dose (6 mg/kg/day) comparison (126 genes), with no significant DEGs in vehicle vs. the highest dose and only five DEGs when PCBs were modeled as a continuous variable.

**Table 1.** Number of differentially expressed genes (DEG) by pairwise PCB dose comparisons or PCB dose as a continuous variable.

|  |  |  |  |  |
| --- | --- | --- | --- | --- |
| **Tissue** | **Sex** | **Comparison** | **Number of DEGs (FDR<0.05)** | **Number of Dose-Response DEGs (FDR<0.05)** |
| Brain | Females | Vehicle vs. 0.1 mg/kg/d | 0 | 5 |
|  |  | Vehicle vs. 1 mg/kg/d | 0 |  |
|  |  | Vehicle vs. 6 mg/kg/d | 0 |  |
|  |  | 0.1 mg/kg/d vs. 1mg/kg/d | 11 |  |
|  |  | 0.1 mg/kg/d vs. 6 mg/kg/d | 126 |  |
|  |  | 1 mg/kg/d vs. 6 mg/kg/d | 1 |  |
|  | Males | Vehicle vs. 0.1 mg/kg/d | 0 | 368 |
|  |  | Vehicle vs. 1 mg/kg/d | 0 |  |
|  |  | Vehicle vs. 6 mg/kg/d | 318 |  |
|  |  | 0.1 mg/kg/d vs. 1mg/kg/d | 11 |  |
|  |  | 0.1 mg/kg/d vs. 6 mg/kg/d | 82 |  |
|  |  | 1 mg/kg/d vs. 6 mg/kg/d | 62 |  |
| Placenta | Females | Vehicle vs. 0.1 mg/kg/d | 190 | 25 |
|  |  | Vehicle vs. 1 mg/kg/d | 37 |  |
|  |  | Vehicle vs. 6 mg/kg/d | 178 |  |
|  |  | 0.1 mg/kg/d vs. 1mg/kg/d | 8 |  |
|  |  | 0.1 mg/kg/d vs. 6 mg/kg/d | 124 |  |
|  |  | 1 mg/kg/d vs. 6 mg/kg/d | 2 |  |
|  | Males | Vehicle vs. 0.1 mg/kg/d | 0 | 3 |
|  |  | Vehicle vs. 1 mg/kg/d | 393 |  |
|  |  | Vehicle vs. 6 mg/kg/d | 360 |  |
|  |  | 0.1 mg/kg/d vs. 1mg/kg/d | 233 |  |
|  |  | 0.1 mg/kg/d vs. 6 mg/kg/d | 393 |  |
|  |  | 1 mg/kg/d vs. 6 mg/kg/d | 0 |  |

Similar to brain, differences in the number of placental DEGs were predominantly monotonic in males, but non-monotonic in females. Male placental DEGs were found at both the medium and high dose group versus vehicle (393 and 360, respectively; Table 1), unlike brain. To find overlapping DEGs across tissue type and sex, an upset plot of DEGs (uncorrected p<0.05) shows the number of overlapping DEGs for the dose-responsive model (Figure 1B). While matched tissues showed the largest overlap regardless of sex, surprisingly, male brain shared many more PCB DEGs with female placenta (375 genes) than male placenta (124). Female brain was an outlier in showing the lowest overlap of PCB DEGs with other tissues.

A secondary objective of this study was to investigate whether dietary FA supplementation during development could counteract effects of developmental PCB exposure. Therefore, we modeled the interaction of low dose PCB (0.1 mg/kg/d) and dietary FA supplementation to identify DEGs with a significant effect modification of FA on PCB exposure. Only female placenta had any DEGs with a significant effect modification of FA on PCB (18; Table 2).

**Table 2.** Number of differentially expressed genes (DEG) by pairwise comparisons of low dose PCBs with or without folic acid (FA) supplementation.

|  |  |  |  |  |
| --- | --- | --- | --- | --- |
| Tissue | Sex | Comparison | Number of DEGs (FDR<0.05) | Number of interxn DEGs (FDR<0.05) |
| Brain | Females | Vehicle vs. Vehicle + FA | 0 | 0 |
|  |  | Vehicle vs. 0.1 mg/kg/d | 0 |  |
|  |  | Vehicle vs. 0.1 mg/kg/d + FA | 1 |  |
|  |  | Vehicle + FA vs. 0.1 mg/kg/d | 1 |  |
|  |  | Vehicle + FA vs. 0.1 mg/kg/d + FA | 0 |  |
|  |  | 0.1 mg/kg/d vs. 0.1 mg/kg/d + FA | 9 |  |
|  | Males | Vehicle vs. Vehicle + FA | 0 | 0 |
|  |  | Vehicle vs. 0.1 mg/kg/d | 0 |  |
|  |  | Vehicle vs. 0.1 mg/kg/d + FA | 1 |  |
|  |  | Vehicle + FA vs. 0.1 mg/kg/d | 0 |  |
|  |  | Vehicle + FA vs. 0.1 mg/kg/d + FA | 1 |  |
|  |  | 0.1 mg/kg/d vs. 0.1 mg/kg/d + FA | 4 |  |
| Placenta | Females | Vehicle vs. Vehicle + FA | 18 | 18 |
|  |  | Vehicle vs. 0.1 mg/kg/d | 1 |  |
|  |  | Vehicle vs. 0.1 mg/kg/d + FA | 47 |  |
|  |  | Vehicle + FA vs. 0.1 mg/kg/d | 0 |  |
|  |  | Vehicle + FA vs. 0.1 mg/kg/d + FA | 6 |  |
|  |  | 0.1 mg/kg/d vs. 0.1 mg/kg/d + FA | 15 |  |
|  | Males | Vehicle vs. Vehicle + FA | 0 | 0 |
|  |  | Vehicle vs. 0.1 mg/kg/d | 0 |  |
|  |  | Vehicle vs. 0.1 mg/kg/d + FA | 12 |  |
|  |  | Vehicle + FA vs. 0.1 mg/kg/d | 0 |  |
|  |  | Vehicle + FA vs. 0.1 mg/kg/d + FA | 0 |  |
|  |  | 0.1 mg/kg/d vs. 0.1 mg/kg/d + FA | 0 |  |

### Experimental design of consensus modules within weighted gene correlation network analyses

Because of the complex relationships observed by sex, dosage, and tissue type from the DEG analyses, we decided to use a systems-based approach to comprehensively understand each of these variables on the transcriptional response to prenatal PCB mixture. Weighted gene correlation network analysis (WGCNA) is a useful way to perform pairwise correlations among genes that allows grouping of co-expressed groups of genes into modules.^19^ Each module is then represented by an eigengene value as a reduced representative of expression levels of all the genes within a module. We are then able to examine the expression dynamics of groups of genes and gene pathways associated with each PCB dose and/or FA supplementation, comparing across the brain-placenta axis and across both sexes.

First, we used a sex-consensus co-expression network of gene expression profiles across males and females, then stratified the data using these sex-consensus modules to investigate sex-specific impacts on the brain versus placenta following prenatal exposures to PCBs (0, 0.1, 1, 6 mg/kg/d). Twenty-three consensus modules between males and females were defined for brain, with an overall preservation score of 0.57 (Additional File 1:

Figure S1, Additional File 2: Table S1), while a separate set of 23 consensus modules between males and females were defined for placenta, with an overall preservation score of 0.60 (Additional File 1: Figure S2, Additional File 2: Table S2).

Second, we used a tissue-consensus co-expression network of gene expression profiles across brain and placenta, then stratified the data using these tissue-consensus modules to investigate tissue-specific PCB effects in males versus females. Twenty-one consensus modules between brain and placenta were defined for males, with an overall preservation score of 0.68 (Additional File 1: Figure S3, Additional File 2: Table S3), while a separate set of 18 consensus modules between brain and placenta were defined f or females, with an overall preservation score of 0.75 (Additional File 1: Figure S4, Additional File 2: Table S4).

Lastly, we constructed networks for the same two module types (sex-consensus and tissue-consensus) but only for samples relevant to the FA supplementation experiment (vehicle, vehicle + FA, 0.1 mg/kg/d PCBs, 0.1 mg/kg/d PCBs + FA) (Additional File 1: Figures S5 - S8, Additional File 2: Tables S5 - S8). For each of these analyses, the modules are arbitrarily named by color. Since these color names are reassigned in each separate module set, the same color can represent a different group of genes in different figure panels. Module colors compared left-to-right in figures are always comparing the same module representing the same group of genes (Figures 2-9).

**Figure 2.**
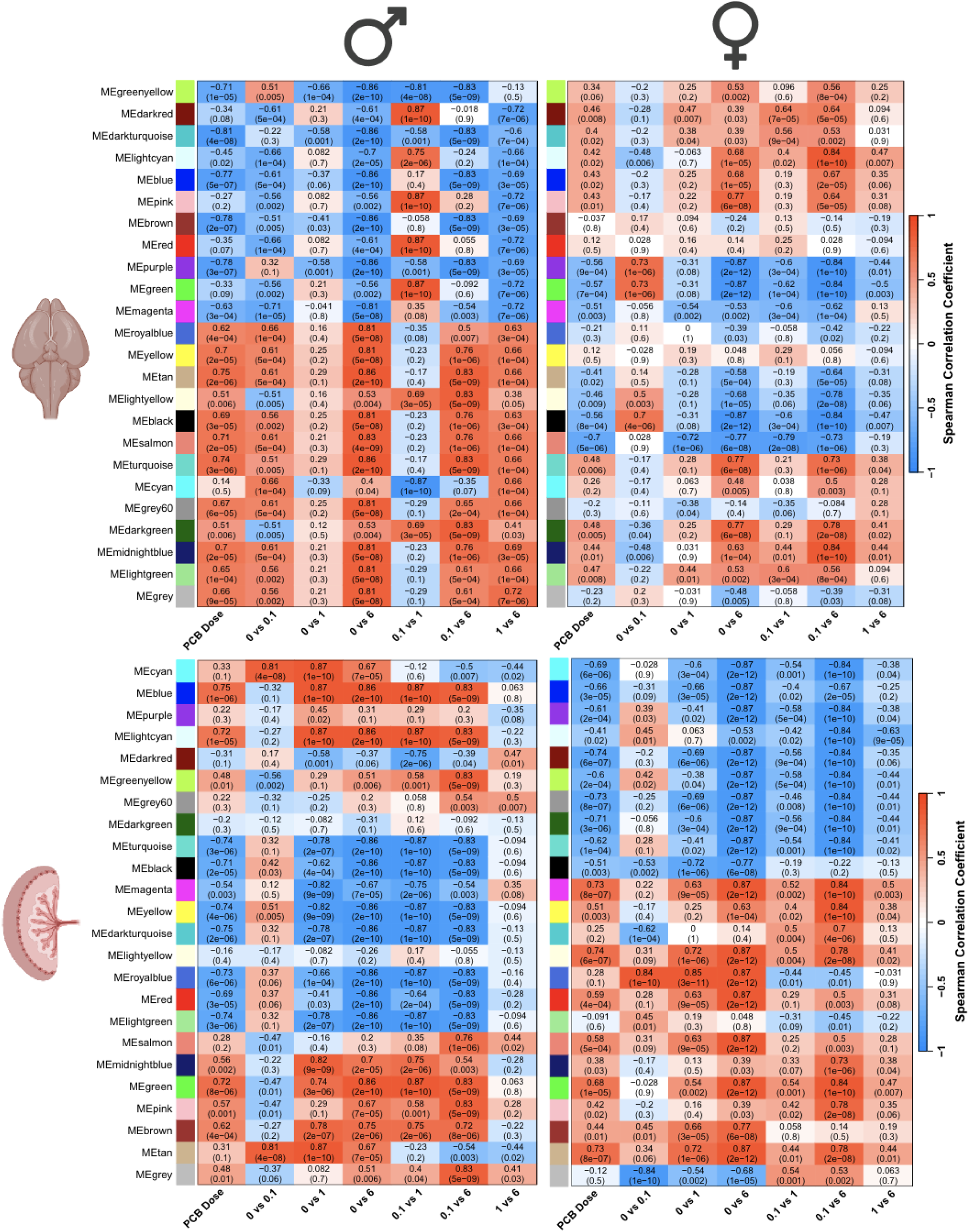
Sex-consensus modules of brain and placenta reveal broad transcriptional impacts with sex-specific and non-monotonic dosage effects in response to prenatal PCB exposure. Two different sets of WGCNA modules (y-axes) were defined for either brain (top) or placenta (bottom) that could be compared across sexes (left-right comparison of the same module). Spearman correlations with each PCB dose comparison (x - axes) are color coded (red, positive; blue, negative) with both r of correlation and p-value provided for each module and trait.

### Sex-consensus modules of brain and placenta reveal broad transcriptional impacts with sex-specific and non-monotonic dosage effects in response to prenatal PCB exposure

The sex-consensus modules allow the same groups of correlated genes to be compared between males and females separately in brain and placenta (Figure 2, y-axes). The module eigengene values, reflecting expression levels of all co-expressed genes within each module, were then correlated with PCB dosage effects, with most modules showing significant (p<0.05) and strong (>0.5 r) positive (red) or negative (blue) associations with PCBs in at least one sex, tissue, or dosage. Considering PCB as a continuous variable (column 1 of x-axes), the male brain samples showed much stronger correlations across all modules than the female brain, a result that was not observed in placenta. Interestingly, several modules are strongly correlated with PCB dose in the same direction between males and females (ie, MEturquoise in brain), while other modules are strongly correlated in the opposite direction between males and females (ie, MEblue in brain). Pairwise comparisons of dosage groups were further investigated to identify potential non-monotonic dosage effects. The most apparent overall non-monotonic effect was the dampened effect in vehicle (0) versus PCB 1 mg/kg/day that was observed across most modules in the male brain, but not female brain or male placenta. These results indicate that the 0.1 and 6 mg/kg/day doses had a greater transcriptomic effect than the 1 mg/kg/day dose, but also that this non-monotonic PCB effect was dependent on both sex and tissue.

For each of the sex consensus modules in brain and placenta, genes in each module were investigated for enrichment within known Gene Ontology (GO) and KEGG pathways. To examine potential gene pathways involved in the sex-specific PCB effects, we focused on modules with significantly enriched KEGG pathway terms that are transcriptionally impacted by PCBs either in the same or different correlation directions between the sexes (Figure 3). Ribosome was the most frequent KEGG term that was predominantly similarly impacted by PCBs in brain and placenta of both sexes. Interestingly, ribosome functions were also enriched in the salmon module in brain, which showed opposite correlations with PCB dose in male versus female brain, but included a distinct group of genes more specifically involved in mitochondrial translation by GO biological process (Supplemental Table 1). The citrate (TCA) pathway was highly enriched in the Greenyellow module in brain that also was correlated with PCB in opposite directions between the sexes. In placenta, sex differences were observed in modules enriched for Hippo and PPAR signaling, as well as Glyoxylate and dicarboxylate and Glycerolipid metabolism.

**Figure 3.**
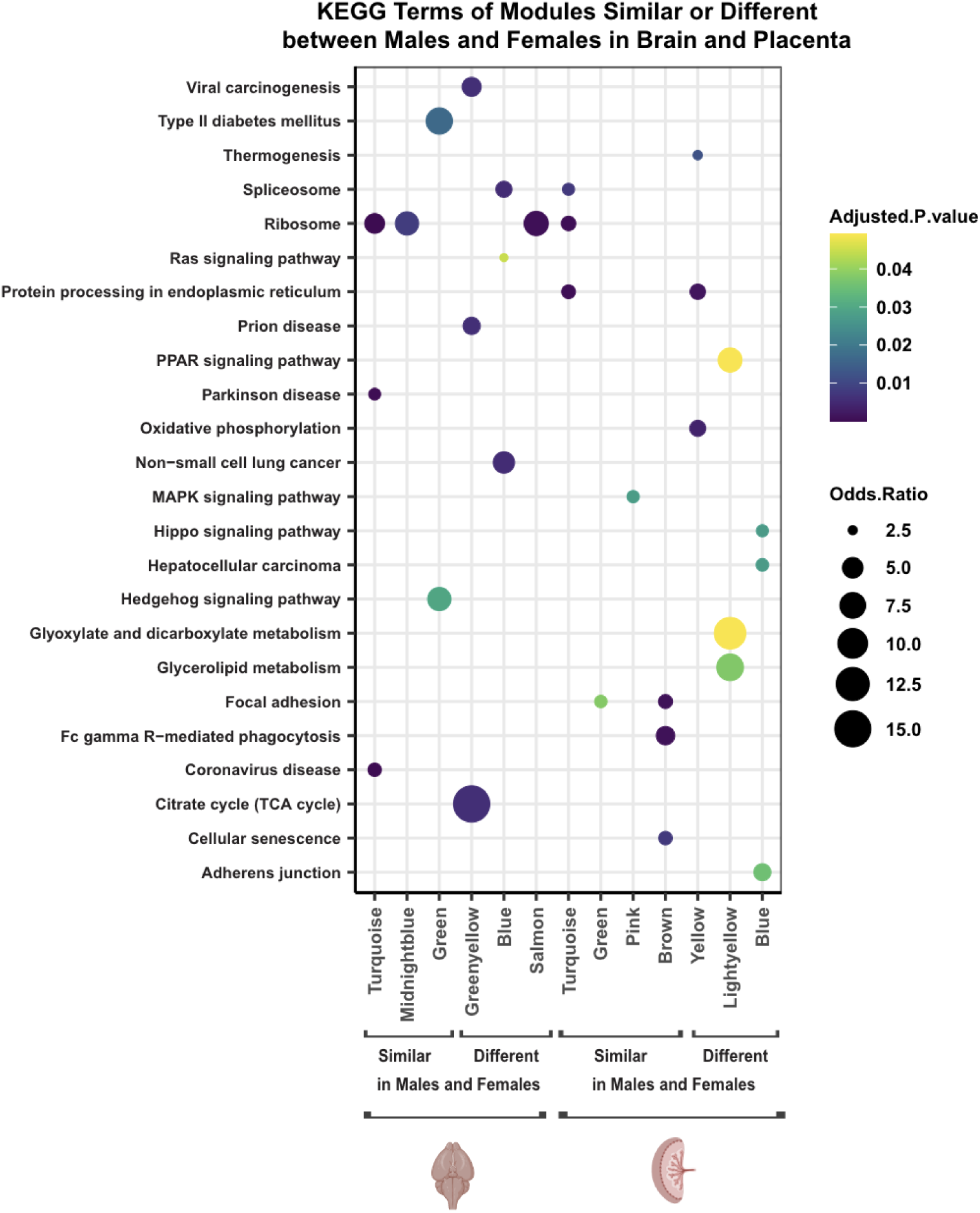
Significantly enriched KEGG gene pathways for sex-consensus modules with significant PCB associations in similar vs. different directions between males and females. Modules that had correlations in the same direction between sexes in at least 5 columns in the Fig. 2 heatmap were called as similar between the sexes. Modules with correlations in the opposite direction between sexes in at least 5 columns were called as different between sexes. The top 3 KEGG terms enriched in these modules, along with its p-values and log of odds ratio are shown (FDR adjusted p-value of 0.05 is considered statistically significant).

### Tissue-consensus modules reveal placenta-brain axis transcriptional differences between sexes enriched in ribosome and oxidative phosphorylation pathways

The tissue-consensus modules allow the comparison of the same groups of correlated genes to be compared between brain and placental separately in males and females (Figure 4, y-axes). Considering PCB as a continuous variable (column 1 of x-axes), the male brain samples showed much stronger correlations across all modules than female brain, a result that was not observed in placenta, and consistent with the results from the sex-consensus modules (Figure 2) and DGE analyses (Figure 1B). While several modules are strongly correlated with PCB dose in the same direction between brain and placenta (ie, MEgreen in males), other modules are strongly correlated in the opposite direction between brain and placenta (ie, MEbrown in males). Non-monotonic effects from pairwise analyses of PCB dosage were detected in most modules in brain, but not placenta, and these effects were strongest in male brain. The pairwise comparisons with the 1 mg/kg/day dose showed reduced correlations with module transcript levels specifically in brain, similar to what was observed using sex-consensus modules. These results further confirm that non-monotonic transcriptional effects of PCBs were sex- and tissue-dependent.

**Figure 4.**
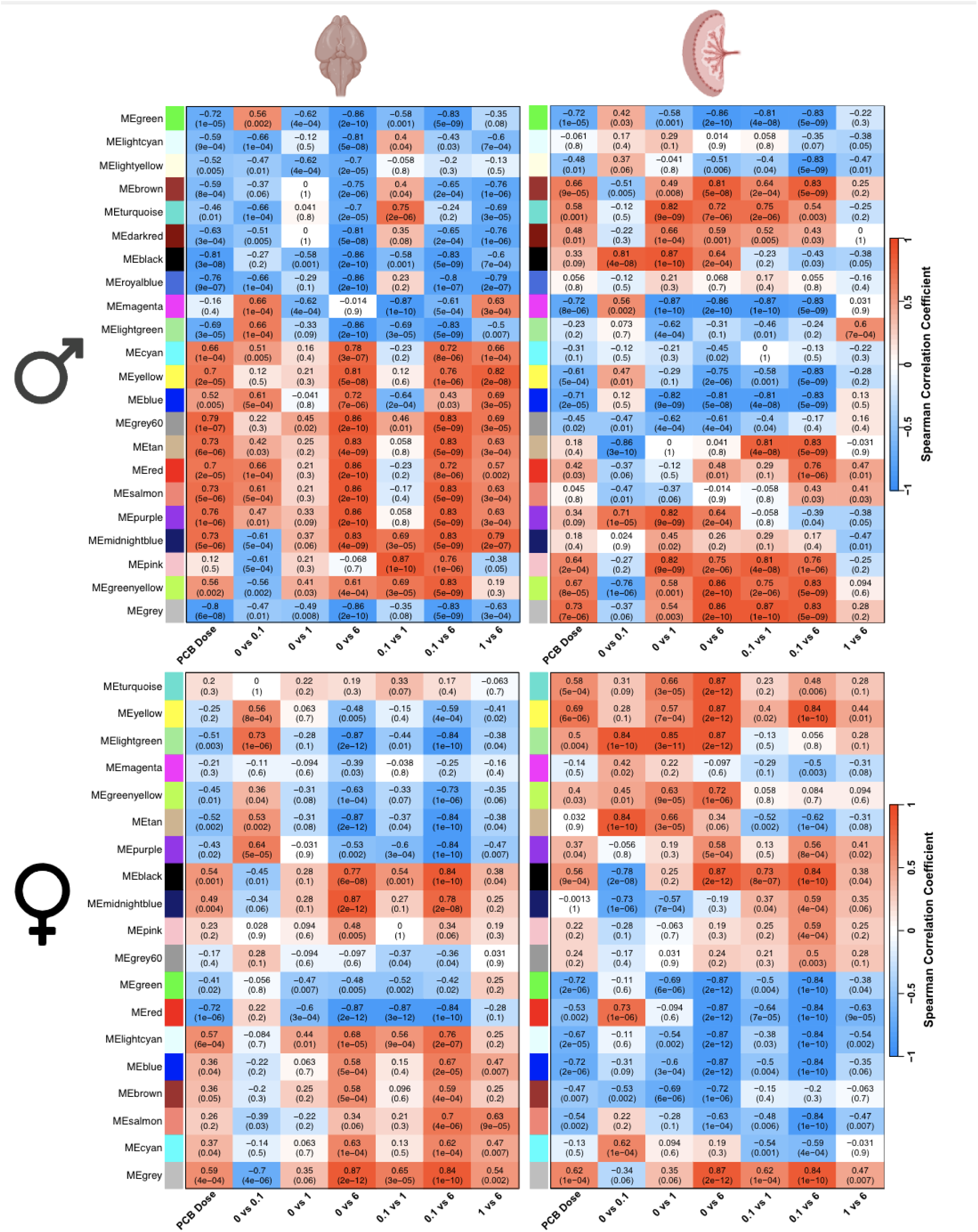
Tissue-consensus modules of brain and placenta reveal broad transcriptional impacts with sex-specific effects in response to prenatal PCB exposure. Two different sets of WGCNA modules (y-axes) were defined for either male (top) or female (bottom) that could be compared across brain and placenta (left-right comparison of same module). Spearman correlations with each PCB dose comparison (x-axes) are color coded (red, positive; blue, negative) with both r of correlation and p value provided for each module and trait.

To examine potential gene pathways involved in the tissue-specific PCB effects across both sexes, we focused on the modules with significantly enriched KEGG pathway terms that are transcriptionally impacted by PCBs either in the same or different correlation directions between brain and placenta (Figure 5). Similar to what was observed from sex-consensus modules, enriched KEGG terms were related to broad pathways in cellular metabolism and the diseases associated with their dysregulation. Interestingly, Ribosome and Oxidative Phosphorylation terms were uniquely enriched in females for two modules (blue and brown) that directionally differed in their transcriptional response between brain and placenta, with a specific downregulation in placenta. The Proteosome KEGG pathway (enriched in magenta in male, green in female) was similarly altered by PCBs in brain and placenta across both sexes, but the TCA pathway was uniquely enriched in a female module (red in female) with similar brain-placenta transcriptional changes. The cyan module in males, enriched for Glutothione and Cytochrome P450 metabolism, was significantly increased in brain (0.1 and 6 mg/kg/d doses) but unchanged (0.1 and 1 mg/kg/d doses) or significantly decreased (6 mg/kg/d) in placenta. The results suggest that the female placenta utilizes different metabolic responses to PCBs involving reduction of energy pathways, while the male placenta shows a lower response to toxins than the male brain.

**Figure 5.**
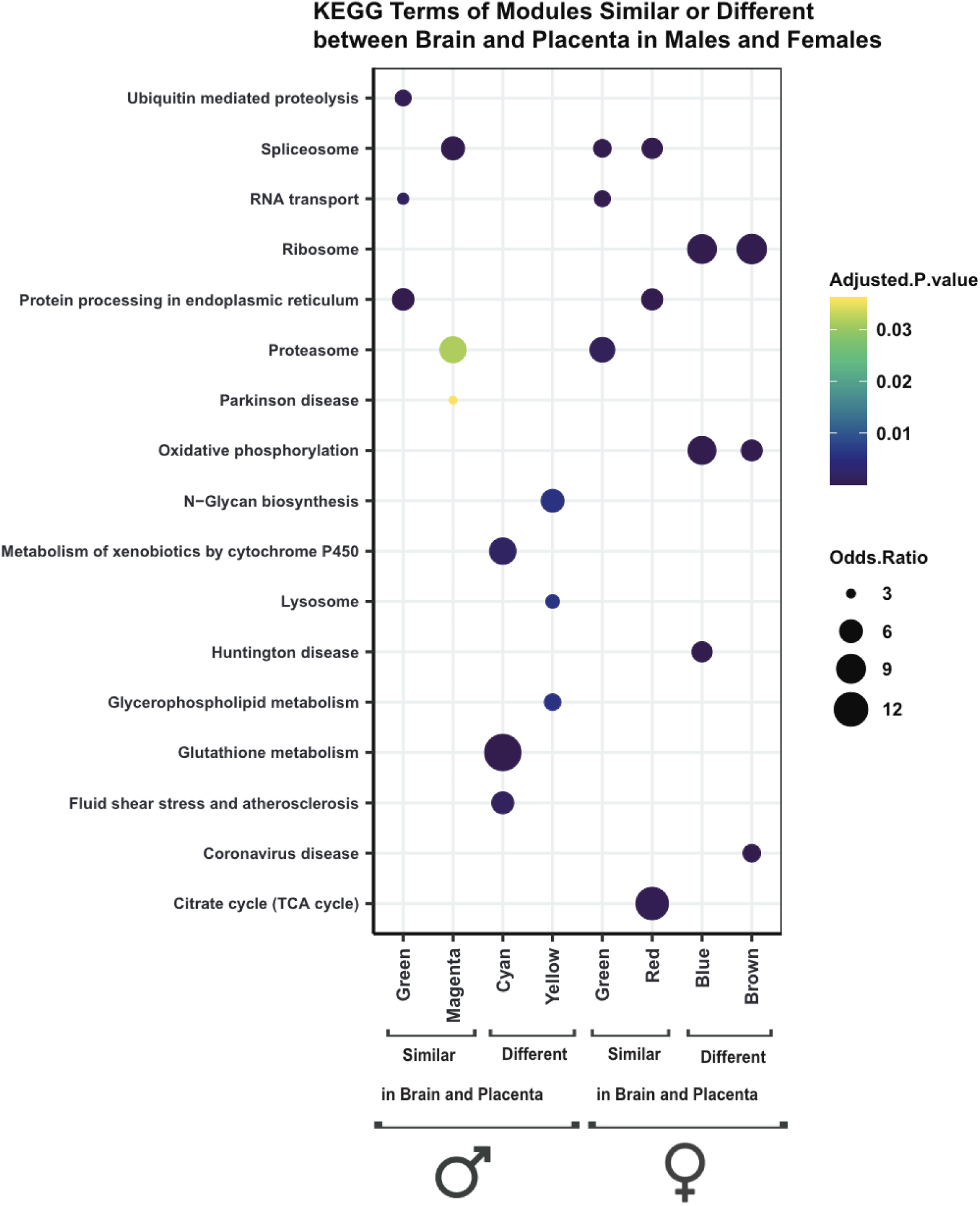
Significantly enriched KEGG gene pathways for tissue-consensus modules with significant PCB associations in similar vs. different directions between brain and placenta. Modules that had correlations in the same direction between tissues in at least 5 columns in the Fig. 4 heatmap were called as similar between tissues. Modules with correlations in the opposite direction between tissues in at least 5 columns were called as different between tissues. The top 3 KEGG terms enriched in these modules, along with its p-values and log of odds ratio are shown (FDR adjusted p-value of 0.05 is considered statistically significant).

### The low-dose PCB transcriptome effects were partially counteracted by folic acid dietary supplementation

To examine potential interaction effects between low dose PCB and dietary folic acid, we performed sex - consensus and tissue-consensus modules for only four groups (0 PCB & No FA, 0.1 mg/kg/d PCB & No FA, 0 PCB & Yes FA, 0.1 mg/kg/d PCB & Yes FA). The heatmaps of all pairwise comparisons for module correlations (Supplementary Figures 5-6) showed that the strongest evidence for interaction effects were for the comparisons of the PCB effect (0 PCB & No FA versus 0.1 PCB & No FA) and the PCB + FA effect (0.1 PCB & No FA versus 0.1 PCB & Yes FA). These two comparisons are shown as heatmaps in Figure 6 (sex-consensus FA) and Figure 7 (tissue-consensus FA) with the modules enriched for significant KEGG terms shown in Figure 8.

**Figure 6.**
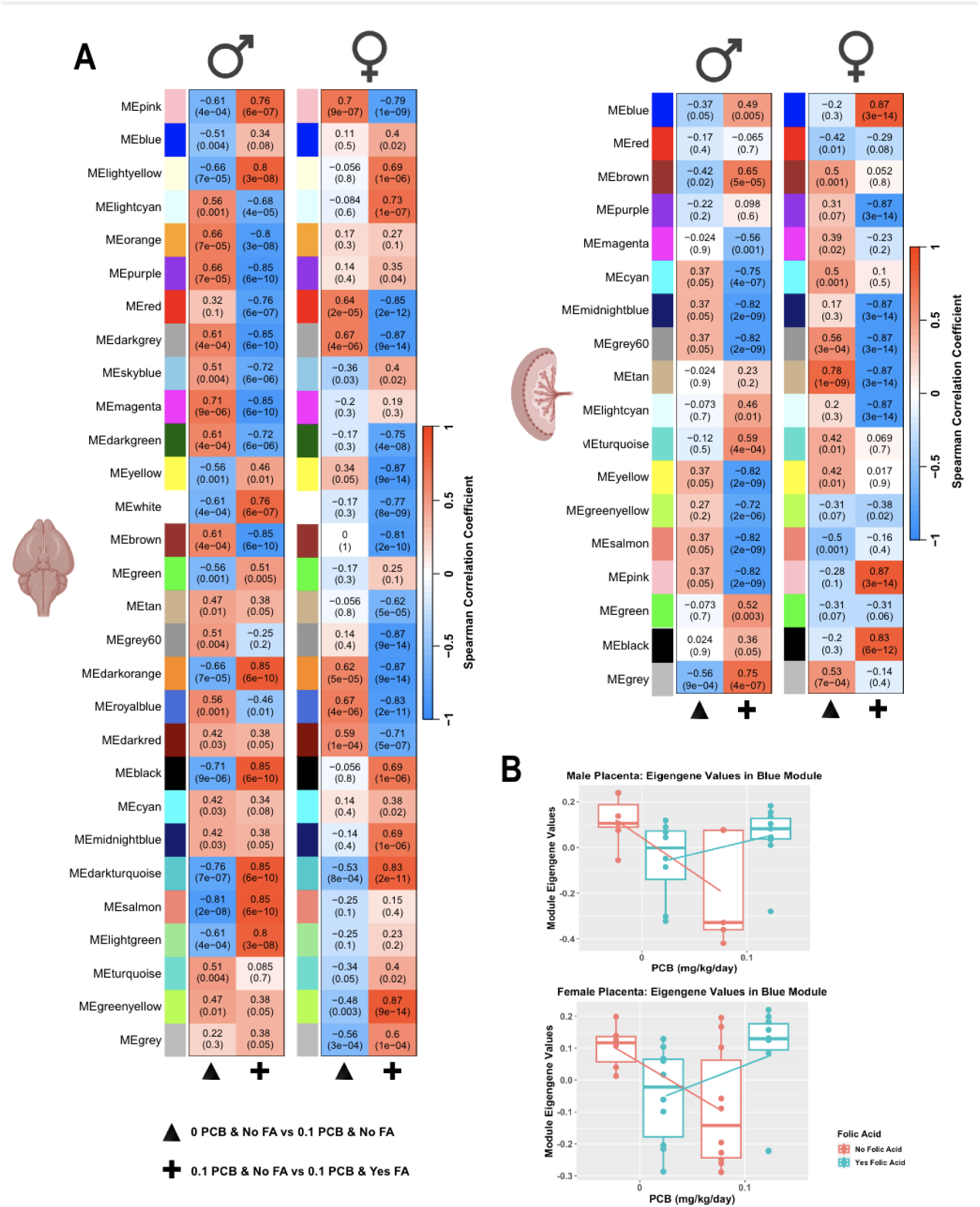
Sex-consensus modules of brain and placenta reveal broad transcriptional impacts with sex-specific counteracting effects of folic acid supplementation on prenatal PCB exposure. **A.** Two different sets of WGCNA modules (y-axes) were defined for either brain (left) or placenta (right) that could be compared across sexes (left-right sex comparison of same module). Only low dose PCB effect without (triangle) or with (+) folic acid (FA) are shown here; additional pairwise comparisons are in Additional File 1: Figure S9. Spearman correlations with each PCB dose comparison (x-axes) are color coded (red, positive; blue, negative) with both r of correlation and p value provided for each module and trait. **B.** Two examples of significant reversal effects are plotted for the Blue module that showed the same effects in male and female placenta.

**Figure 7.**
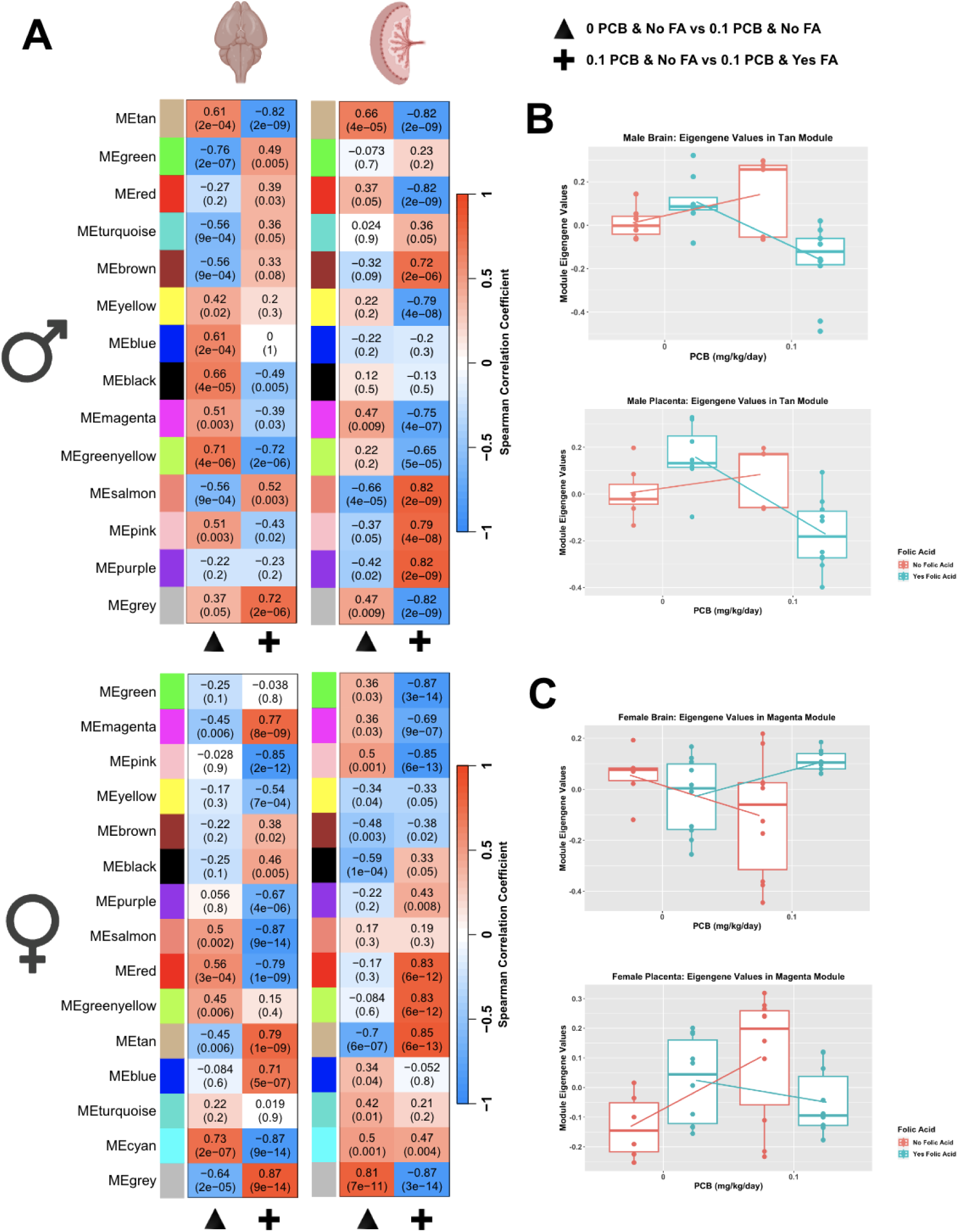
Tissue-consensus modules of brain and placenta reveal broad transcriptional impacts with sex-specific counteracting effects of folic acid supplementation on prenatal PCB exposure. **A.** Two different sets of WGCNA modules (y-axes) were defined for either male (top) or female (bottom) that could be compared across sexes (left-right sex comparison of same module). Only low dose PCB effect without (triangle) or with (+) folic acid (FA) are shown here; additional pairwise comparisons are in Additional File 1: Figure S10. Spearman correlations with each PCB dose comparison (x-axes) are color coded (red, positive; blue, negative) with both r of correlation and p value provided for each module and trait. **B.** Examples of significant reversal effects are plotted for the Tan module that showed the same effects in placenta and brain. **C**. Examples of opposite FA effects are plotted for the Magenta module.

**Figure 8.**
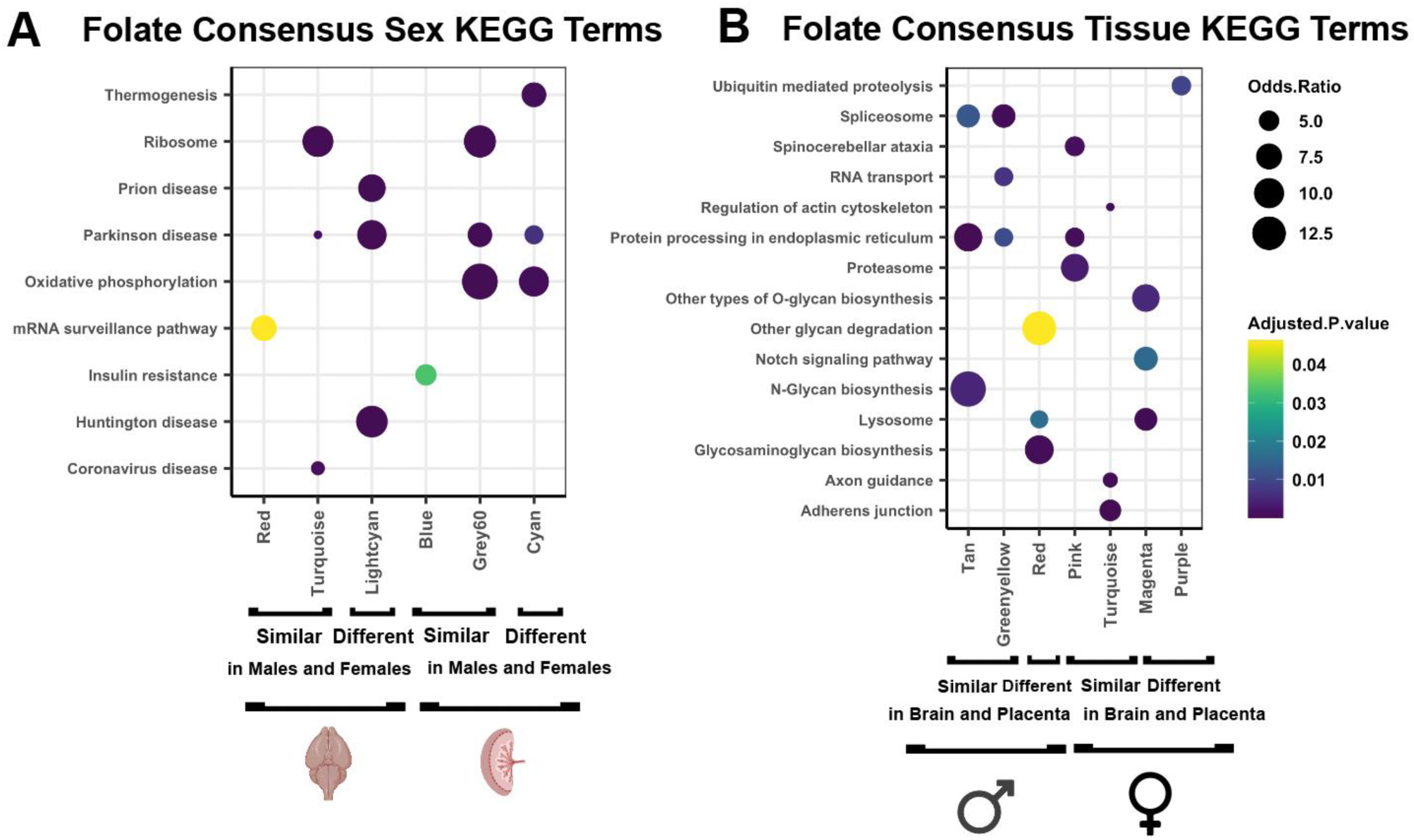
Significantly enriched KEGG gene pathways for folic acid + low dose PCB sex- and tissue-consensus modules. Modules that had correlations in the same direction between sex or tissues in at least 5 columns in the Fig. 6-7 heatmaps were called as similar between tissues. Modules with correlations in the opposite direction between tissues in at least 5 columns were called as different between tissues. The top 3 KEGG terms enriched in these modules, along with its p-values and log of odds ratio are shown (FDR adjusted p-value of 0.05 is considered statistically significant).

The PCB-FA sex-consensus modules showed the expected strong associations with low dose PCB alone (column 1, triangle) in male brain (Figure 6A) as well as strong associations in the opposite direction in the presence of FA. Female brain also showed opposite directions of association with FA+PCB even though the PCB effects alone were less prominent than those in male brain. In placenta, there were modules that were similar and others that were different between male and female, and an opposite effect of FA+PCB in modules of both categories (Figure 6A). The interaction effects of low dose PCB and FA are plotted for individual placental samples for the Blue sex-consensus module to show an example of FA’s counteracting effect on the PCB effect of lowering transcript levels (Figure 6B).

The PCB-FA tissue-consensus modules also showed a counteracting effect of FA in multiple modules, although in general the module associations with FA+PCB effects were stronger in the male placenta than brain (Figure 7A). An example is plotted for the PCB and FA interaction effect in brain versus placenta for the tan module (Figure 7B). The FA+PCB interaction effect in females showed stronger tissue differences than in males, as female transcriptomes showed six modules with opposite directional FA+PCB associations between brain and placenta, compared to only three in males (Figure 7A). For example, the magenta tissue-consensus module in females showed increased transcript levels in placenta but decreased levels in brain in response to PCBs, but both effects were counteracted by FA supplementation (Figure 7C). Also, the tan module in females contained *Xist* as well as its enhancer *Ftx* and *Ogt*, all transcripts known to be higher in females.

KEGG pathway enrichment analyses for PCB-FA sex-consensus modules revealed similar pathways that were observed for PCB dosage effects, including Ribosome, Oxidative phosphorylation, and diseases associated with these pathways, with the Thermogenesis pathway being more enriched in modules showing sex differences (Figure 8A). The FA tissue-consensus modules that were different between brain and placenta were uniquely enriched in Lysosome (male and female), Other glycan degradation (male only), Other types of O-glycan biosynthesis (female only), and Notch signaling (female only) (Figure 8B). Together, these results demonstrate that the counteracting effect of FA on PCB-induced transcriptional changes is acting broadly through energy - regulating gene pathways, with some differences unique to female placenta.

## Discussion

This study was designed to understand molecular pathways in the brain-placenta axis that may explain the male bias in NDD risk, utilizing an existing mouse model of prenatal exposure to a human-relevant mixture of PCBs. Transcriptomic analyses of embryonic brain and matched placenta were performed using system biology-based approaches, resulting in several novel findings. First, prenatal PCB exposure resulted in widespread dysregulation of correlated gene networks enriched in energy and cellular metabolism, but the direction of these effects varied between males and females, as well as between brain and placenta. Second, the female-specific placental transcriptional responses uniquely enriched in citrate metabolism, O-glycan biosynthesis, and decreased energy consumption corresponded to relatively muted transcriptional responses in female compared to male brain. Third, there was a strong non-monotonic effect of PCB dosage, with the lowest and highest dosages showing the strongest transcriptional responses, but this effect was both sex- and tissue-dependent, and most prominent in male brain. Lastly, folic acid supplementation was able to reverse most of the gene networks’ transcriptional changes, but these interaction effects were most prominent in female placenta.

Together, these results provide evidence for a protective effect of the female placenta in modulating the expression of metabolic pathways in a way that reduces the transcriptional impact on the developing brain. These results also have translational implications for the use of human placental tissue in the prediction of molecular changes in brain as well as the prevention of NDDs with wider use of prenatal vitamins containing folic acid.

The neurotoxicity of PCBs is predicted to be related to its known effects on calcium channels and neuronal synapses,^20–21^ but genome-wide analyses of molecular changes associated with PCB exposures are scant. Our previous whole-genome analyses of DNA methylation changes resulting from a 1 mg/kg/d dose of the same PCB mixture and mouse model used in this study is consistent with the strong overlap in dysregulated gene pathways between placenta and brain, even though the direction of change was sometimes directionally opposite.^18^ In the current study, we directly examined sex and tissue differences as a potential explanation for the directionally different responses of brain versus placenta. Since our study was the first to our knowledge to employ WGCNA approach to bioinformatic analyses of PCB effects, we were able to uniquely demonstrate broad changes across multiple gene networks enriched for functions in energy and cellular metabolism that were distinctly altered in females compared to males. Our results are consistent with a prior transcriptome and lipidome study in zebrafish brain that showed disruption of energy homeostasis could be explained by impaired pathways of mitochondrial function and lipid metabolism regulation following exposure to an environmentally -relevant mixture of PCBs and PBDEs.^22^ Our results showing enriched mitochondrial and oxidative phosphorylation gene pathways are also consistent with a transcriptomic analysis of newborn mouse dentate granule cells following perinatal exposure to Aroclor 1254, a commercial PCB mixture.^23^ What is unique about our investigation is the integration of placenta and fetal brain and the demonstration of profound sex differences in the transcriptional response to prenatal PCB exposure in both brain and placenta.

Sex differences related to PCB exposures have been previously observed, but poorly understood.^17, 24^ Sex differences in response to prenatal PCB exposures have been mixed in the human epidemiological studies examining secondary sex ratio.^25,26^ However, a recent large population study found multiple air and water pollutants associated with changes to the sex ratio at birth, including PCBs which associated with male bias^27^. In a recent analysis of six different prenatal chemical mixtures with metabolic syndrome (MetS) score stratified by sex, PCB mixture was uniquely associated with a higher MetS score in female children and lower MetSscore in male children, which is consistent with our findings of large-scale sex differences in transcriptional modulation of metabolic genes in placenta. While differences in sex hormones are usually implicated in transcriptional sex differences, females also differ from males genetically by the presence of two X chromosomes and the lack of a Y chromosome. A prior study demonstrated that the X-linked gene *O*-GlcNAc transferase (OGT) gene has higher expression in female placenta and placental OGT was required for female protective effects of prenatal stress on mitochondrial energy production.^28^ Interestingly, we also observed an enrichment for O-glycan biosynthesis gene pathways that work upstream of OGT in protective female modules responding to PCB combined with folic acid protection. Notch signaling pathways were also enriched in females, while glycan degradation pathways that degrade Notch-related proteins were enriched in males perinatally exposed to PCBs. Together, these results are consistent with sex differences in metabolic pathway activation in the placenta-brain axis that may be encoded in the imbalance in OGT and other X- or Y-linked genes with differential expression, such as *Xist* and *Ftx*. While our results do not rule out the influence of sex hormones, no significant KEGG pathways were identified to explain the large-scale PCB-induced transcriptional differences between the sexes in placenta and brain. These results together with the sex-differential DNA methylation PCB effects we observed previously in the placental-brain axis of this PCB exposure model suggest that wide-spread epigenetic differences on metabolic pathways are primarily due to the genetic differences in sex chromosome content between the sexes.

In the current study, we expanded the PCB mixture doses to a total of three (0.1, 1, and 6 mg/kg/d) and discovered a strong non-monotonic dose response, with the medium dose evoking the lowest transcriptional response. Our results showing that the non-monotonic response to *in utero* exposure to PCBs was dependent on both sex and tissue is consistent with our prior studies of behavioral and neuronal dendritic phenotypes in the same PCB mixture mouse model.^14^ Non-monotonic responses have been frequently described previously for PCB and other endocrine disrupting chemicals, but are poorly understood. Of the six different molecular mechanisms described to explain non-monotonic effects,^29^ our results are most consistent with a dose-dependent metabolism modulation^30^ because of the sex-specificity and large number of genes and gene pathways regulating metabolism in the placental-brain axis.

While PCB levels are gradually declining in the environment and human bodies over recent decades, our finding of broadly dysregulated fetal brain transcription combined with our prior finding of social behavioral impairments in males at the 0.1 mg/kg/d dose suggests that PCB neurotoxicity may not be declining with environmental exposures. What was potentially promising from our results is that dietary folic acid supplementation was successful in reducing the transcriptional impacts of PCBs by acting on similarly disrupted metabolic pathways, including oxidative phosphorylation and O-glycan pathways. Prenatal vitamins containing folic acid are recommended for all women planning pregnancy to prevent neural tube defects and potentially other neurodevelopmental conditions like autism spectrum disorder, but only ∼50% of women in the MARBLES cohort took prenatal vitamins in the first month of pregnancy or before which is the window of protection in humans. Our results suggest the importance of public health education campaigns about the importance of taking prenatal vitamins prior to conception to prevent possible harms from low-level PCBs and other pollutants.

Our results also have implications for early detection of NDDs at birth. As the major source of nutrients and other support to the fetus, placenta is a tissue source rich in epigenetic and transcriptional biomarkers that can be indicative of the *in utero* environment. Since placenta is a readily available tissue source discarded at birth, it can be used to assess NDD risk by diagnostic screening approaches at birth. Outcomes of this project can potentially lead to lowering the age at which children are diagnosed with ASD and other NDDs, which can improve the effectiveness of pharmacological and behavioral interventions during the most plastic window of postnatal brain development. Our findings are also expected to have clinical relevance beyond NDDs. Enriched KEGG pathways for PCB-altered genes identified the neurodegenerative disorders Parkinson and Huntington disorders, as well as Coronovirus and Prion disease, all of which may have oxidative phosphorylation and metabolic pathways in their pathogenesis. A prior study of PCB-180 effects on human iPSC-derived neurons also identified Parkinson disease pathway genes.^31^ Since sex differences are apparent in prevalence and/or severity of both neurodevelopmental and neurodegenerative disorders, our results highlight the importance of the placental-brain axis for understanding sex by environmental interactions in brain health.

## Methods

### Mouse exposure model and tissue harvest

The PCB MARBLES mixture, consisting of PCB 28 (48.2%), PCB 11 (24.3%), PCB 118 (4.9%), PCB 101 (4.5%), PCB 52 (4.5%), PCB 153 (3.1%), PCB 180 (2.8%), PCB 149 (2.1%), PCB 138 (1.7%), PCB 84 (1.5%), PCB 135 (1.3%) and PCB 95 (1.2%), was formulated to mimic the 12 most abundant congeners identified from the serum of pregnant women in the ASD-enriched MARBLES cohort, as previously described.^32^ C57BL/6J female mice (Jackson Laboratory) aged 6 to 8 weeks were orally exposed to 4 different doses (0, 0.1, 1.0, or 6.0 mg/kg/d) of the PCB mixture in peanut oil (Spectrum Organic) and peanut butter (Trader Joe’s Organic Creamy). The PCB mixture was stored as concentrated stock in peanut oil, then diluted in peanut butter to make dosing mixes for the four dose groups (0, 0.1, 1, and 6 mg/kg/d). The dosing mixes were premade for the entire experiment and stored in the refrigerator and remixed daily. For alterations to folic acid in the chow, mice were fed ad libitum, with the AIN-93G purified diet, containing with or without 6 ppm folic acid supplementation (ENVIGO 190757, 94045, respectively). Females were dosed daily for at least 2 weeks before breeding commenced. Daily dosing then continued through breeding and gestation, up to embryonic day 18 (E18). On E18, pregnant females were euthanized in a CO_2_ euthanasia chamber. Immediately after euthanasia, embryos were removed, and tissues dissected and flash frozen. Embryo and tissue weights were recorded, as well as relative position within the uterine horns. Table 3 shows the total number of samples harvested for transcriptomic analyses per treatment group.

**Table 3.** Number of samples harvested from ∼3 dams per group, sexes combined.

|  |  |  |  |
| --- | --- | --- | --- |
| **Dosing Group** | **96 Matched Brain/Placenta Samples** | **Female Samples** | **Male Samples** |
| 0 PCB / 0 FA | 13 Brain<br>13 Placenta | 6 | 7 |
| 0.1 PCB / 0 FA | 15 Brain<br>15 Placenta | 10 | 5 |
| 1 PCB / 0 FA | 15 Brain<br>15 Placenta | 9 | 6 |
| 6 PCB / 0 FA | 15 Brain<br>15 Placenta | 6 | 9 |
| 0 PCB / Yes FA | 18 Brain<br>18 Placenta | 10 | 8 |
| 0.1 PCB / Yes FA | 20 Brain<br>20 Placenta | 10 | 10 |

**Table 4.** Settings for Module generation in WGCNA.

| Co-expression Network | Samples (N = ) | Power | Minimum Genes in Module | Number of Modules Generated |
| --- | --- | --- | --- | --- |
| PCB Sex-Consensus Brain | 27 Male<br>31 Female | 22 | 50 | 24 |
| PCB Sex-Consensus Placenta | 27 Male<br>31 Female | 18 | 150 | 24 |
| PCB Tissue-Consensus Male | 27 Brain<br>27 Placenta | 16 | 100 | 22 |
| PCB Tissue-Consensus Female | 31 Brain<br>31 Placenta | 20 | 100 | 19 |
| FA Sex-Consensus Brain | 28 Male<br>35 Female | 28 | 50 | 29 |
| FA Sex-Consensus Placenta | 30 Male<br>36 Female | 28 | 150 | 18 |
| FA Tissue-Consensus Male | 30 Brain<br>30 Placenta | 28 | 100 | 12 |
| FA Tissue-Consensus Female | 35 Brain<br>36 Placenta | 24 | 100 | 13 |

### RNA Isolation and RNA-sequencing

RNA was isolated from snap frozen placenta and brain tissue by homogenizing tissue using a TissueLyser II (Qiagen) followed by the All-Prep DNA/RNA/miRNA Universal Kit (Qiagen). Isolated RNA underwent quantification and QC using a Bioanalyzer Eukaryotic Total RNA Nano Assay (Agilent). Libraries were constructed using the KAPA mRNA HyperPrep kit (Roche) and NEXTFLEX Unique Dual Index Barcodes (PerkinElmer). Libraries were pooled and sequenced on a NovaSeq 6000 S4 flow cell (Illumina) for 150 bp paired end reads resulting in approximately 25 million uniquely mapped reads per sample.

### Bioinformatic Analyses

Raw RNA-seq fastq files were processed by trimming adapters using Trim Galore (v0.6.5) and alignment to the mouse genome (mm10) followed by gene count quantification using STAR (v2.7.3a).^33^ QC metrics were generated with MultiQC (v1.9).

### Differential Gene Expression Analysis

Differential expression and subsequent pathway enrichment analyses were carried out using R version 4.1.0. Counts were filtered and normalized using limma voom followed by fitting a linear mixed model using dream weights (variancePartition package) to model litter as a random effect. The eBayes() function was used to test for differential expression, and a cutoff of false discovery rate (FDR<0.05) was used to define differentially expressed genes. Three different models were fit: 1) gene expression ∼ group with contrasts fit to compare each group, 2) gene expression ∼ PCB dose as a continuous variable, and 3) gene expression ∼ PCB + Folic Acid + PCB*Folic Acid to examine genes that had a significant effect modification of Folic Acid on PCB dose. ComplexUpset was used to create UpSet plots of gene overlaps across groups. enrichR followed by GO term slimming with rrvgo was used for pathway enrichment analysis.

### Weighted Gene Correlation Network Analysis

Weighted gene correlation networks (Langfelder and Horvath, 2007) were generated for several subsets of the RNA-seq data in order to compare similar modules of highly correlated genes across either sex or tissue, as well as to test the interaction effects of a single PCB dose (0.1 mg/kg/) and folic acid (FA) supplementation. Four separate consensus networks were generated for PCB dosage effects between the sexes (PCB sex-consensus networks for placenta and brain) and PCB dosage effects between the tissues (PCB tissue-consensus networks for males and females). Four separate consensus networks were generated for PCB + FA supplementation effects (Yes/No FA, vehicle PCB mg/kg/day with Yes/No FA 0.1 ppm PCB) between the sexes and between the tissues (PCB+FA sex-consensus networks for placenta and brain) and (PCB+FA tissue-consensus networks for males and females). A matrix of 13,241 gene counts were used in all consensus networks. Hierarchical clustering was performed to identify and remove sample outliers for each consensus network.

Scale-free topology was used to determine the soft-power threshold to which pairwise correlations between genes are raised. A scale-free topology model fit of at least 0.85 was used to determine the soft-power threshold (Supplemental Figures 9-16). Signed networks were used to maintain the direction of co-expression information among genes. Tukey biweight midcorrelation was used for the network construction, which is a robust method to downweight potential outliers. The minimum number of genes per module were then optimized in order to achieve between 10-30 modules of highly correlated genes. Module eigengenes, essentially the first principal component of a module that serves as a reduced representation of the gene expression of all genes within the module, were computed for each module. Spearman correlation was then used to find correlations between module eigengenes (ME) and treatment group comparisons (traits) because the spearman correlation test addresses non-linear module-trait relationships and can address non-monotonic dosage effects.

entrezIDs were converted to gene symbols using the biomart database in R to allow gene ontology analysis and KEGG pathway analysis to be performed for every module within each consensus network. Modules that had correlations in the same direction (between sex or tissue) in at least 5 columns in the heatmap were considered to be acting similarly (between sex or tissue), and correlations in the opposite direction (between sex or tissue) in at least 5 columns were considered to be acting differently (between sex or tissue). The top 3 KEGG terms enriched in these modules, along with its p-values and log of odds ratio are shown (a FDR adjusted p-value of 0.05 is considered statistically significant).

## Acknowledgements

This work was supported by a National Institutes of Health (NIH) grant (R01ES029213) to JML/PJL/RJS, the UC Davis Intellectual and Developmental Disabilities Research Center (IDDRC) (P50HD103526) and the UC Davis Environmental Health Sciences Center (P30ES023513). Kelly Chau was supported by a NIEHS-funded predoctoral fellowship (T32 ES007059). Its contents are solely the responsibility of the authors and do not necessarily represent the official views of the NIEHS or the NIH. The library preparation and sequencing was carried out by the DNA Technologies and Expression Analysis Cores at the UC Davis Genome Center and was supported by a NIH Shared Instrumentation Grant (1S10OD010786-01). The synthesis of the PCB mixture was supported by the Superfund Research Center at The University of Iowa (P42 ES013661).

