## Additional File 1 for "Placental-brain axis in females detected within broadly impacted metabolic gene networks protects against prenatal PCB exposure"

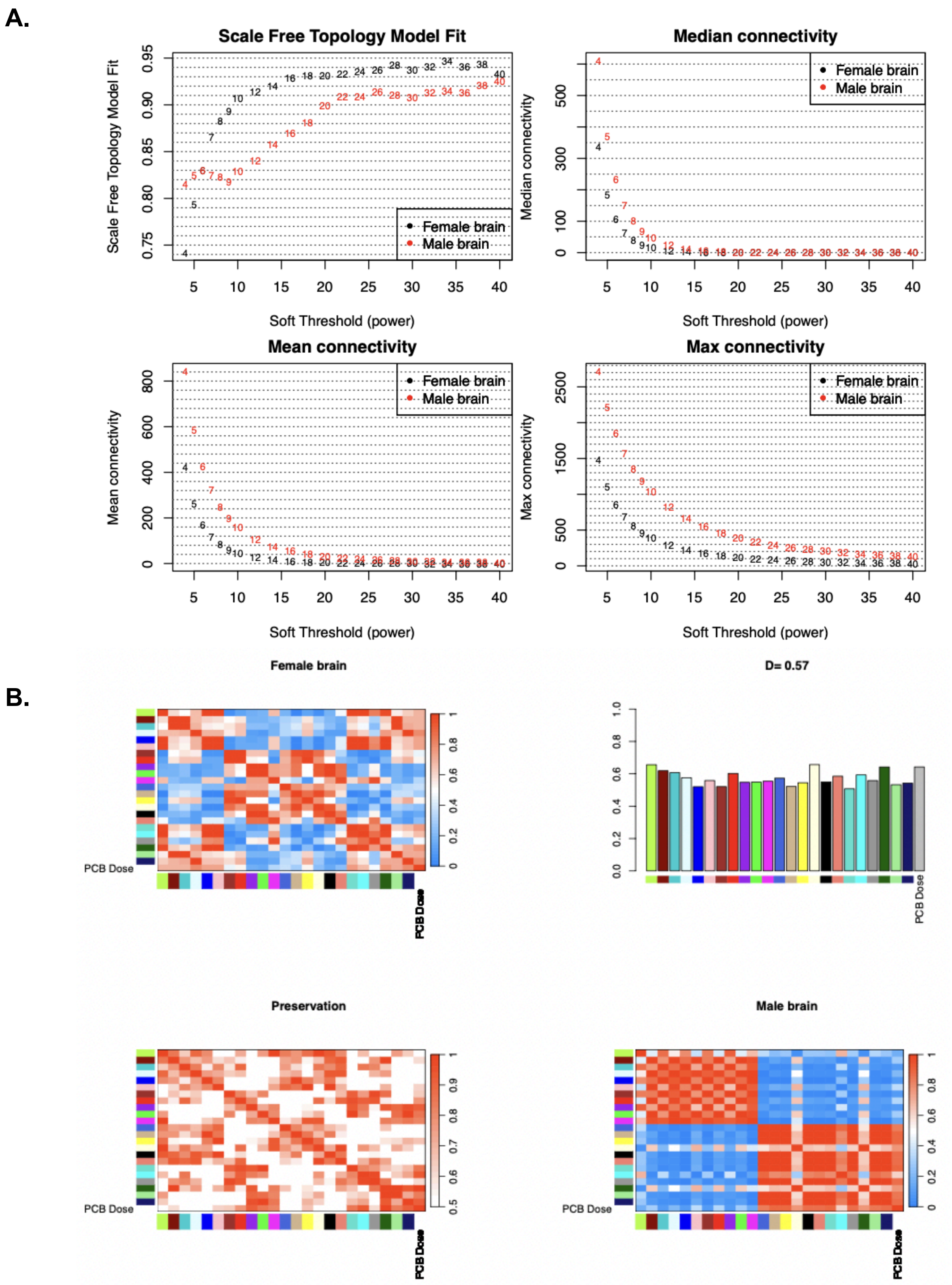


**Figure S1: PCB sex-consensus networks for brain. A)** Network topology analysis. A power of 22 was selected to construct scale-free networks. **B)** Module preservation statistics. The overall preservation of the eigengene networks is denoted by D=0.57.

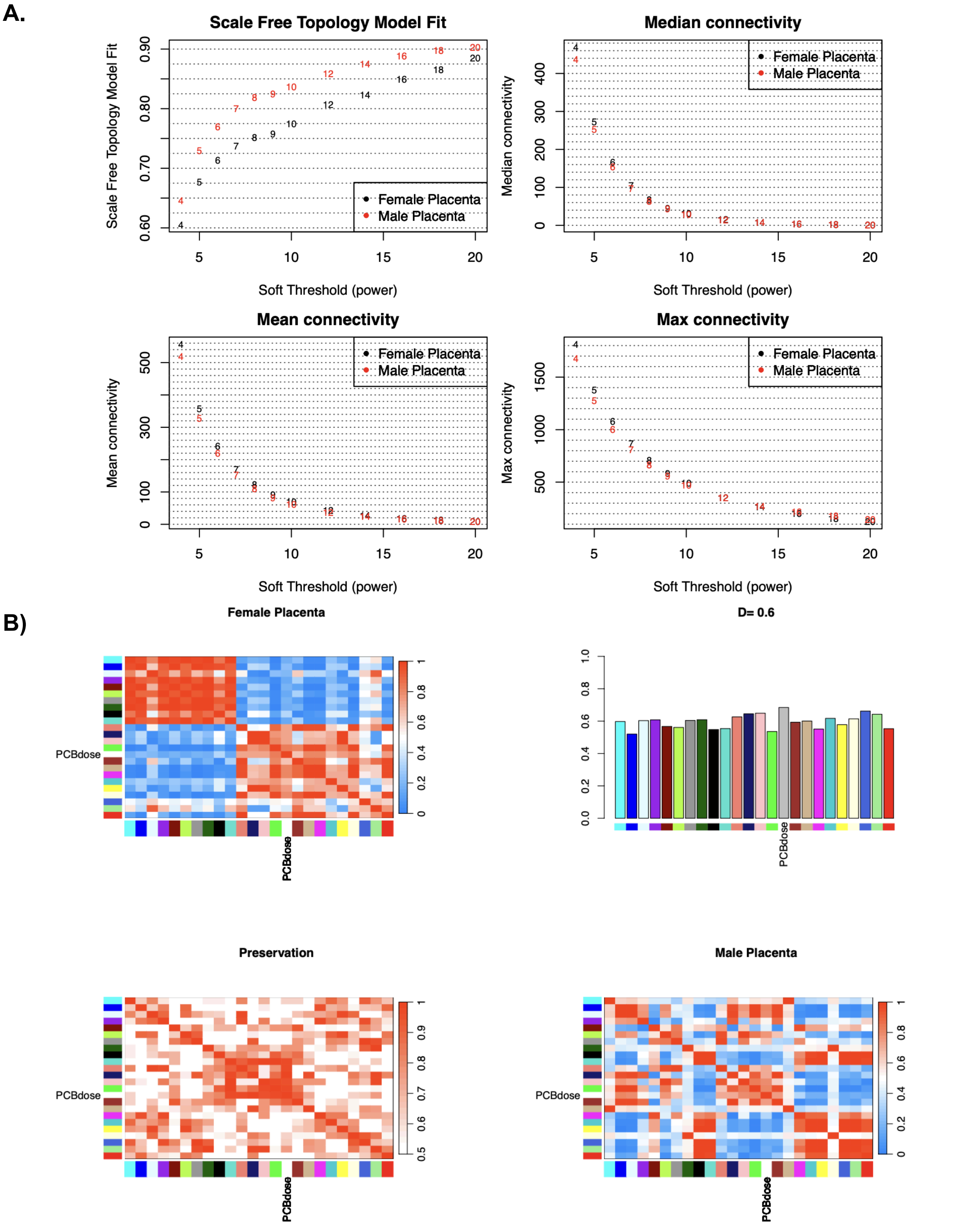


**Figure S2: PCB sex-consensus network for placenta. A)** Network topology analysis. A soft-thresholding power of 18 was selected to construct scale-free networks. **B)** Module preservation statistics. The overall preservation of the eigengene networks is denoted by D=0.6.

A.

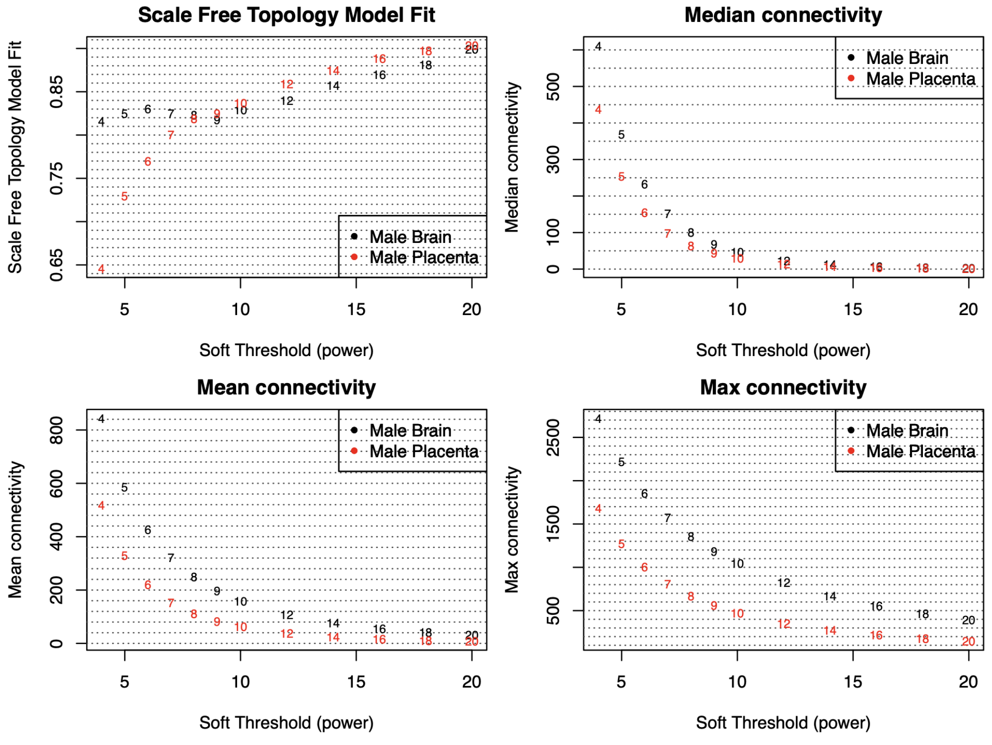


B.

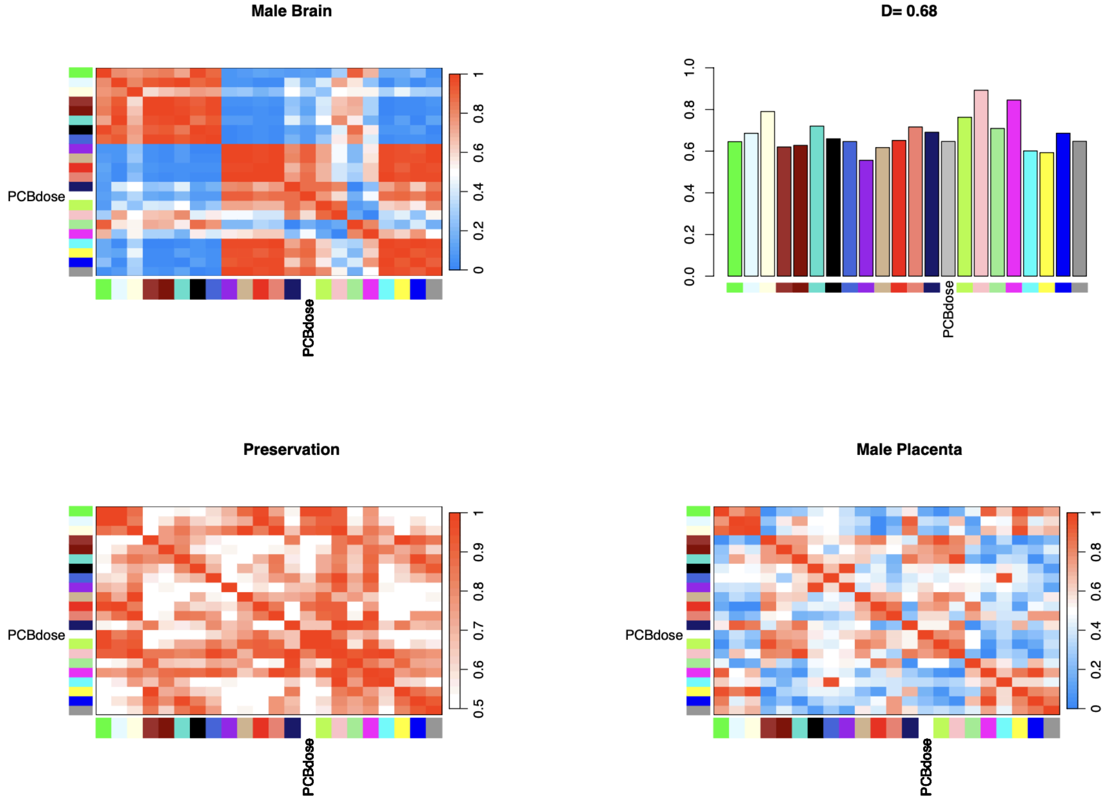


**Figure S3: PCB tissue-consensus networks for males. A)** Network topology analysis. A power of 16 was selected to construct scale-free networks. **B)** Module preservation statistics. The overall preservation of the eigengene networks is denoted by D=0.68.

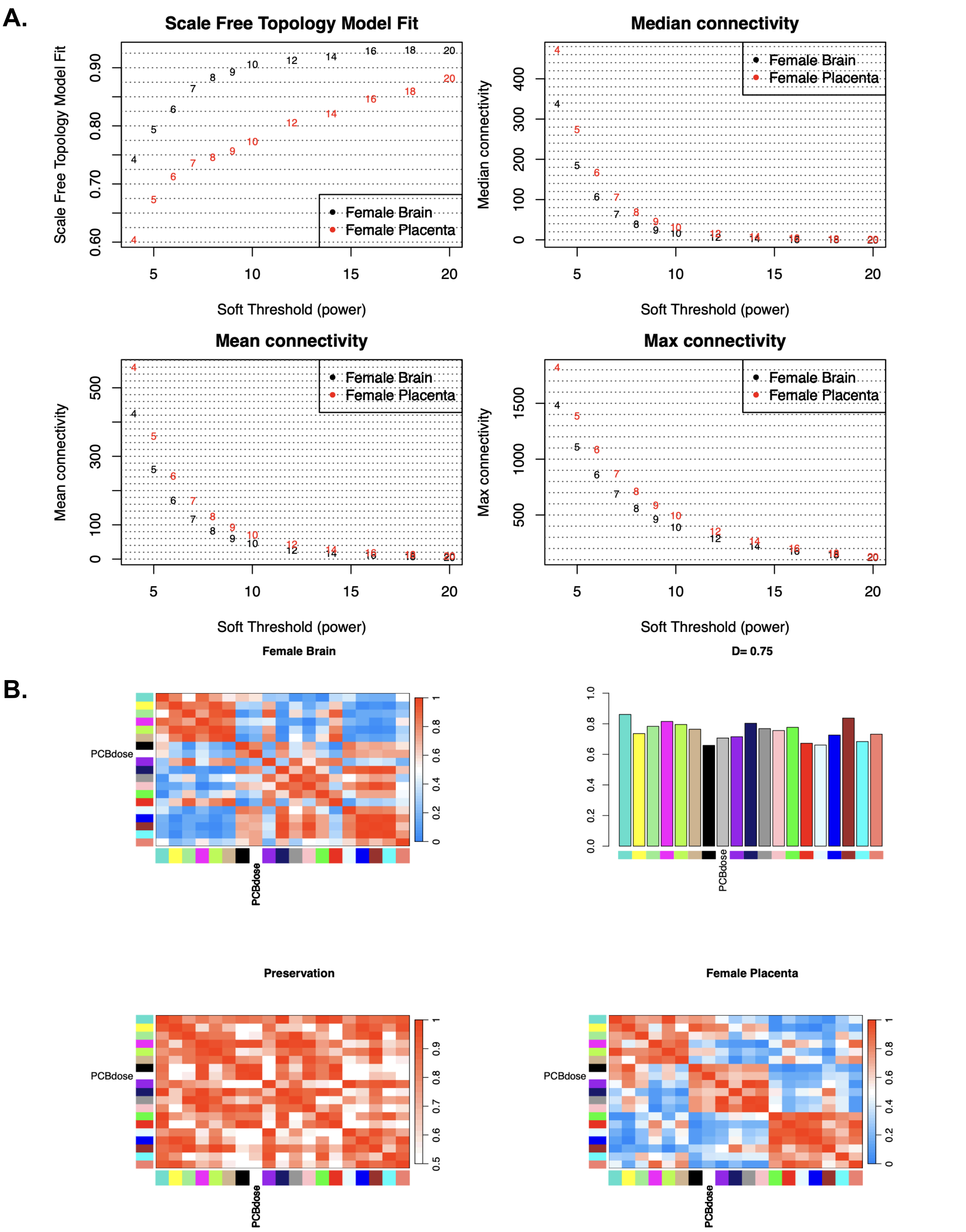


**Figure S4: PCB tissue-consensus networks for females. A)** Network topology analysis. A power of 20 was selected to construct scale-free networks. **B)** Module preservation statistics. The overall preservation of the eigengene networks is denoted by D=0.75.

**A.**

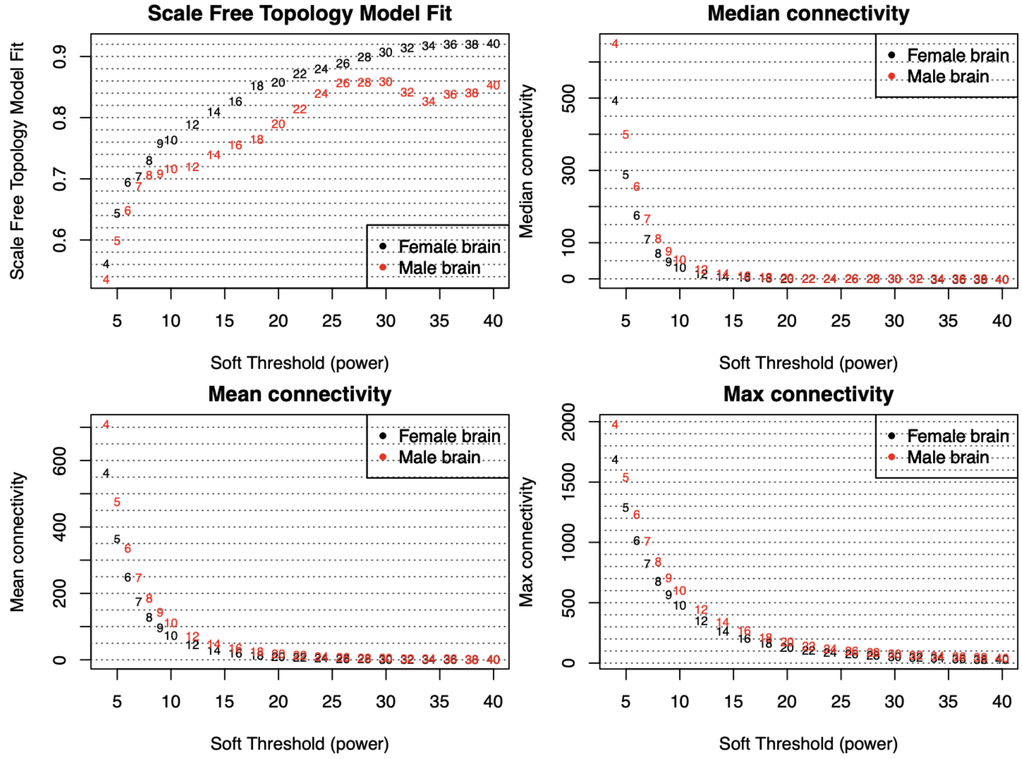


**B.**

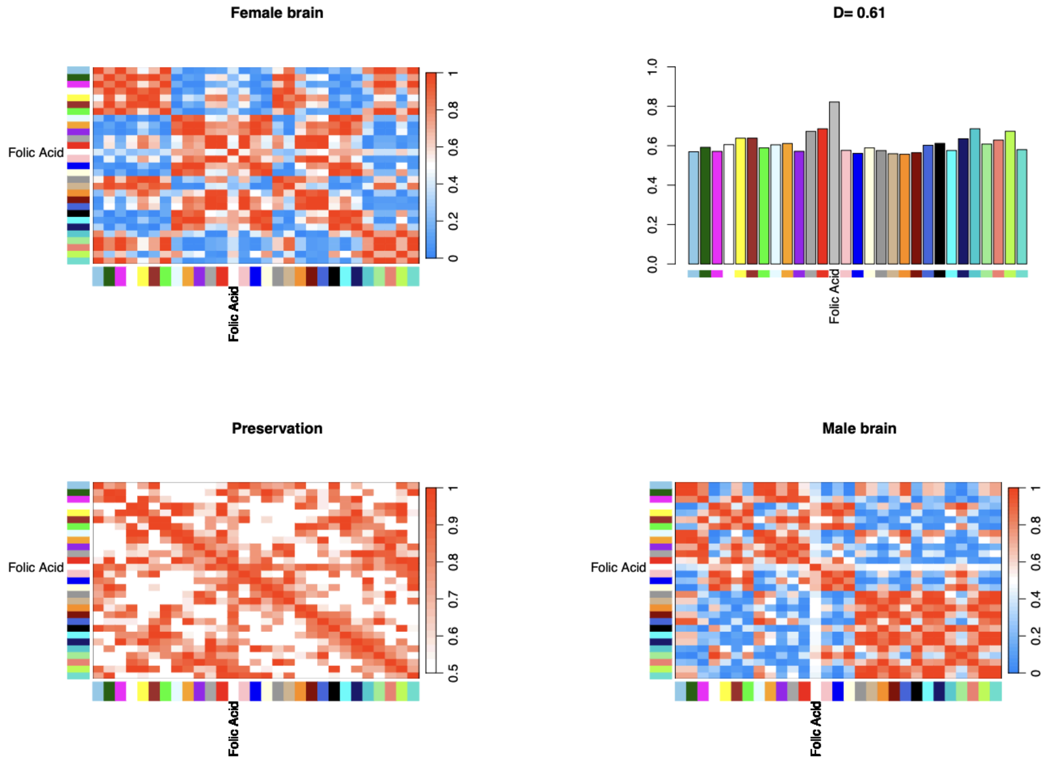


**Figure S5: Folic Acid sex-consensus network for brains. A)** Network topology analysis. A soft-thresholding power of 28 was selected to construct scale-free networks. **B)** Module preservation statistics. The overall preservation of the eigengene networks is denoted by D=0.61.

**A.**

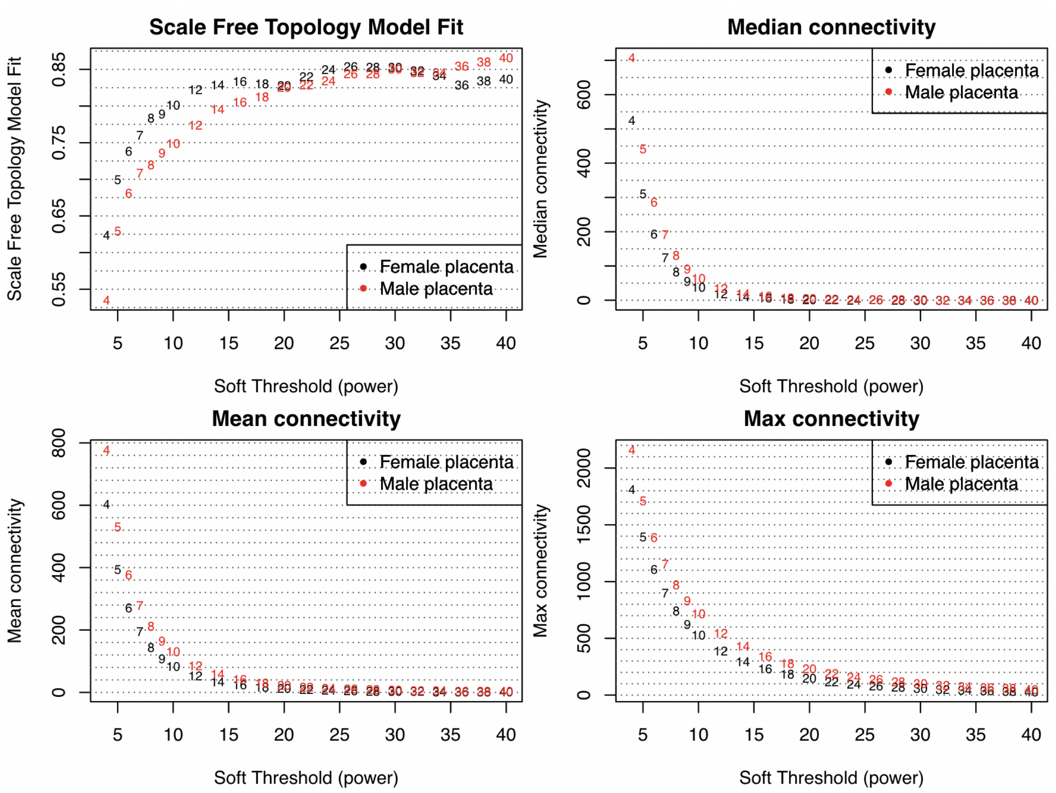


**B.**

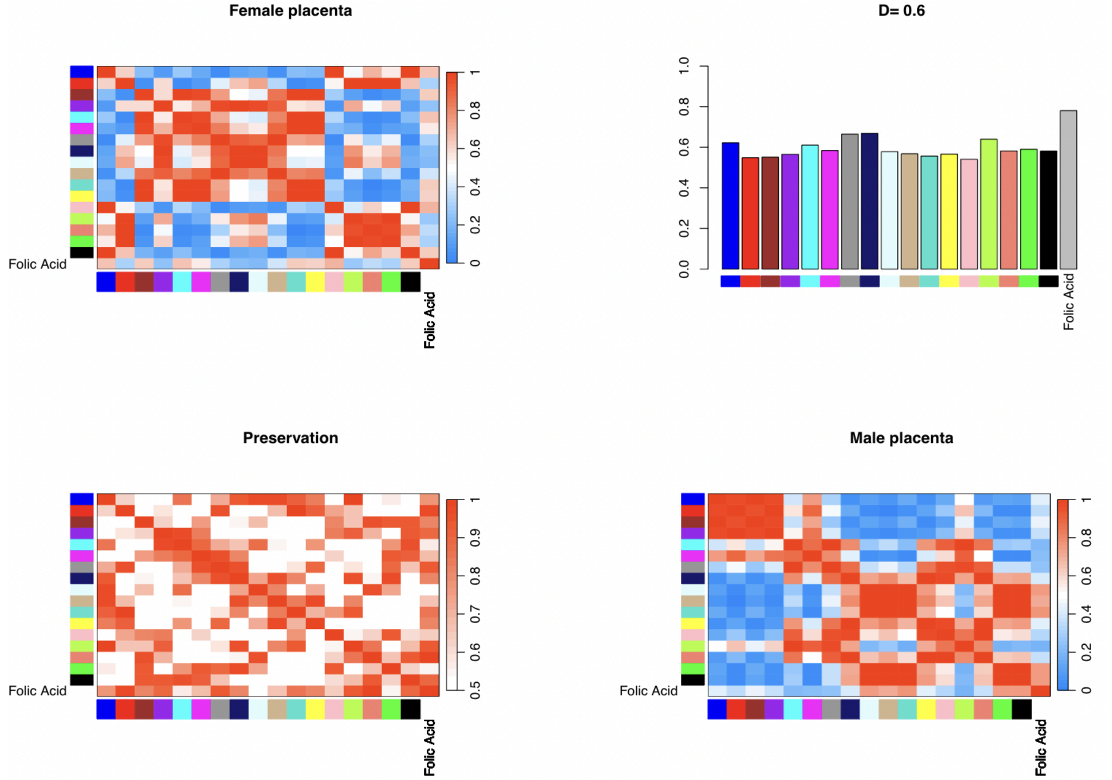


**Figure S6: Folic Acid sex-consensus networks for placenta. A)** Network topology analysis. A soft-thresholding power of 28 was selected to construct scale-free networks **B)** Module preservation statistics. The overall preservation of the eigengene networks is denoted by D=0.6.

A.

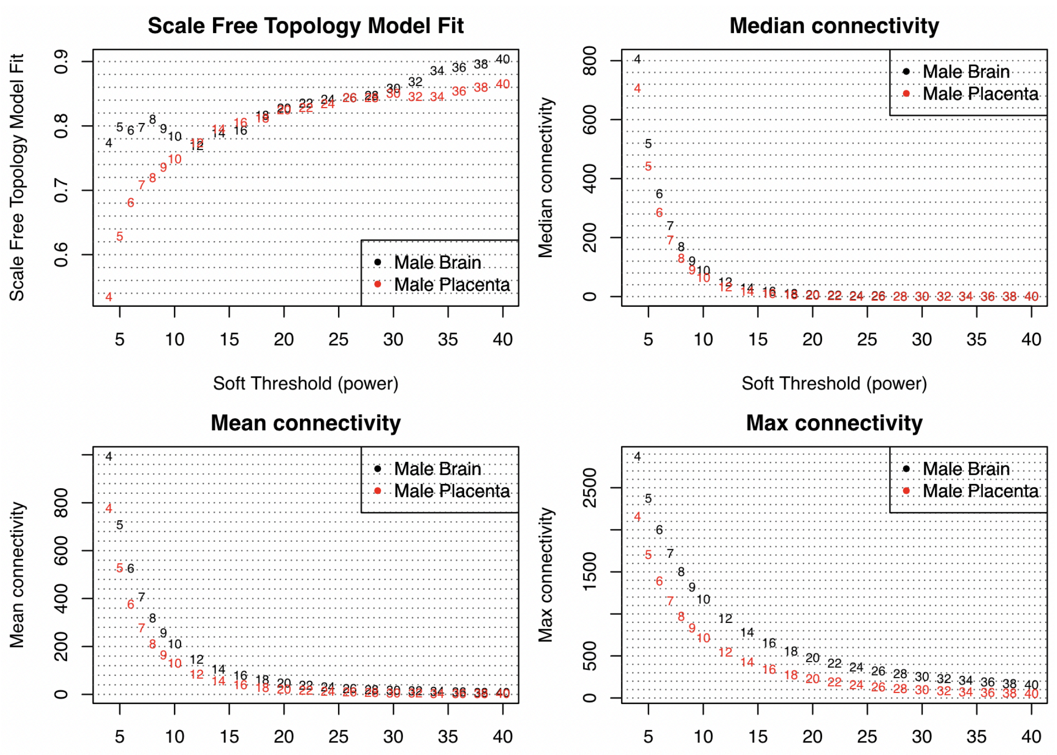


B.

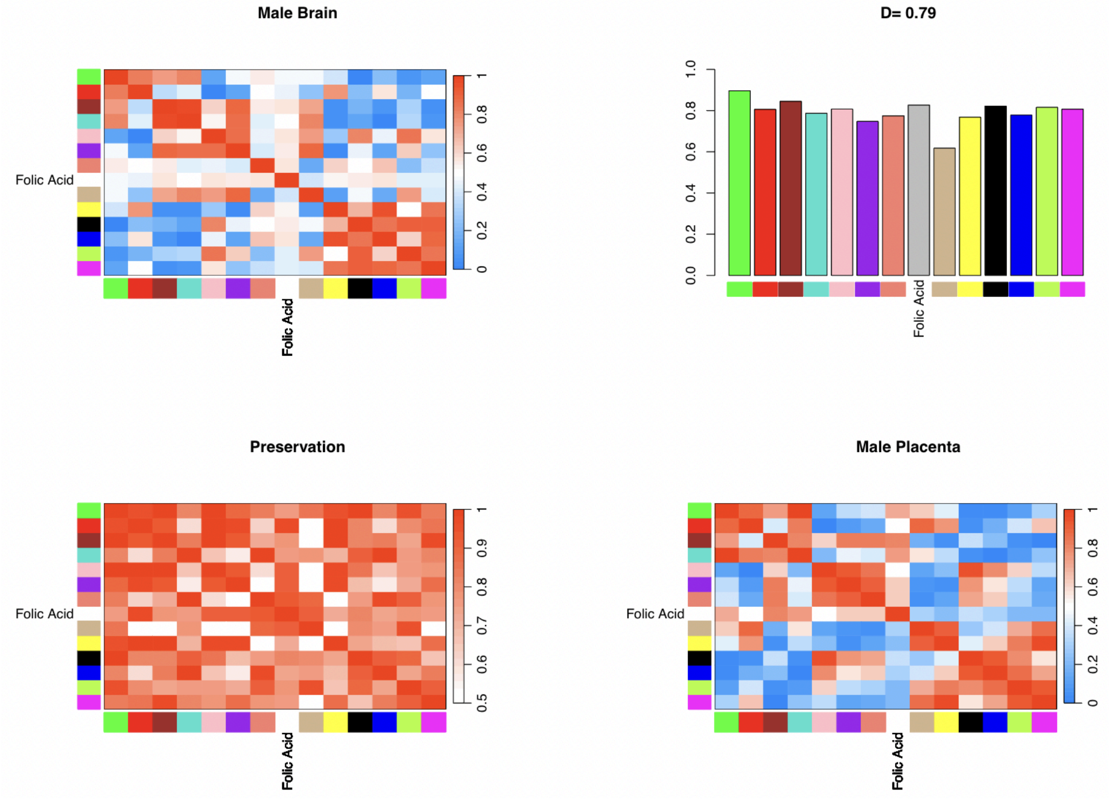


**Figure S7: Folic Acid tissue-consensus networks for males. A)** Network topology analysis. A soft-thresholding power of 28 was selected to construct scale-free networks. **B)** Module preservation statistics. The overall preservation of the eigengene networks is denoted by D=0.79.

A.

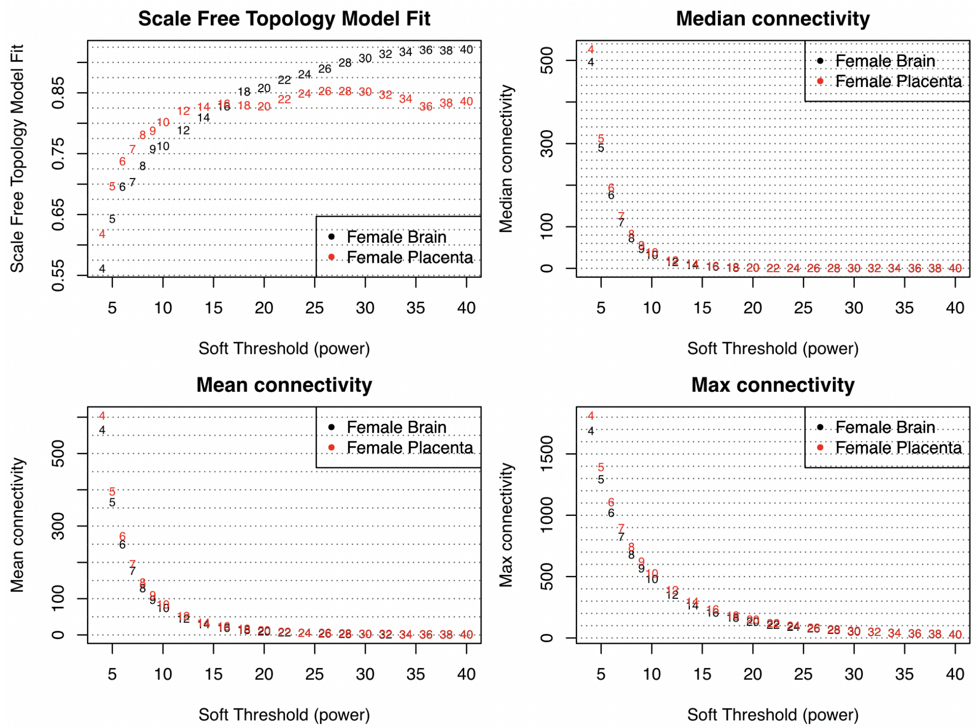


B.

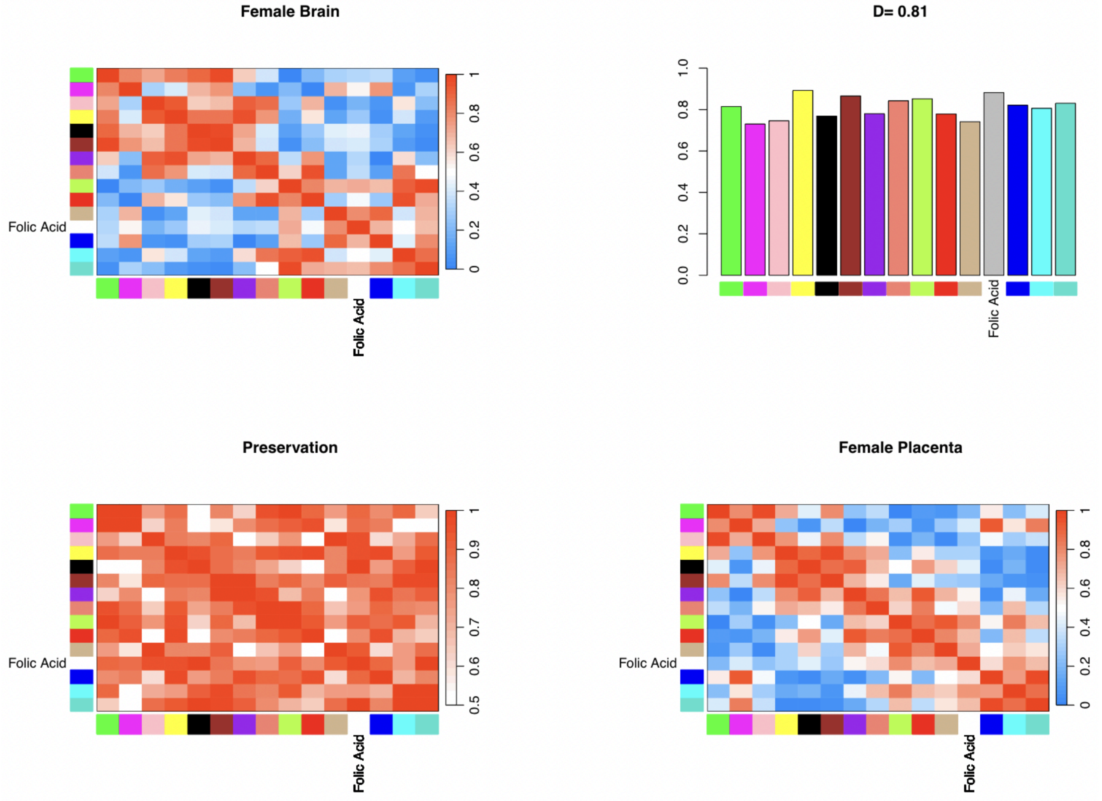


**Figure S8: Folic Acid tissue-consensus networks for females. A)** Network topology analysis. A soft-thresholding power of 24 was selected to construct scale-free networks. **B)** Module preservation statistics. The overall preservation of the eigengene networks is denoted by D=0.81.

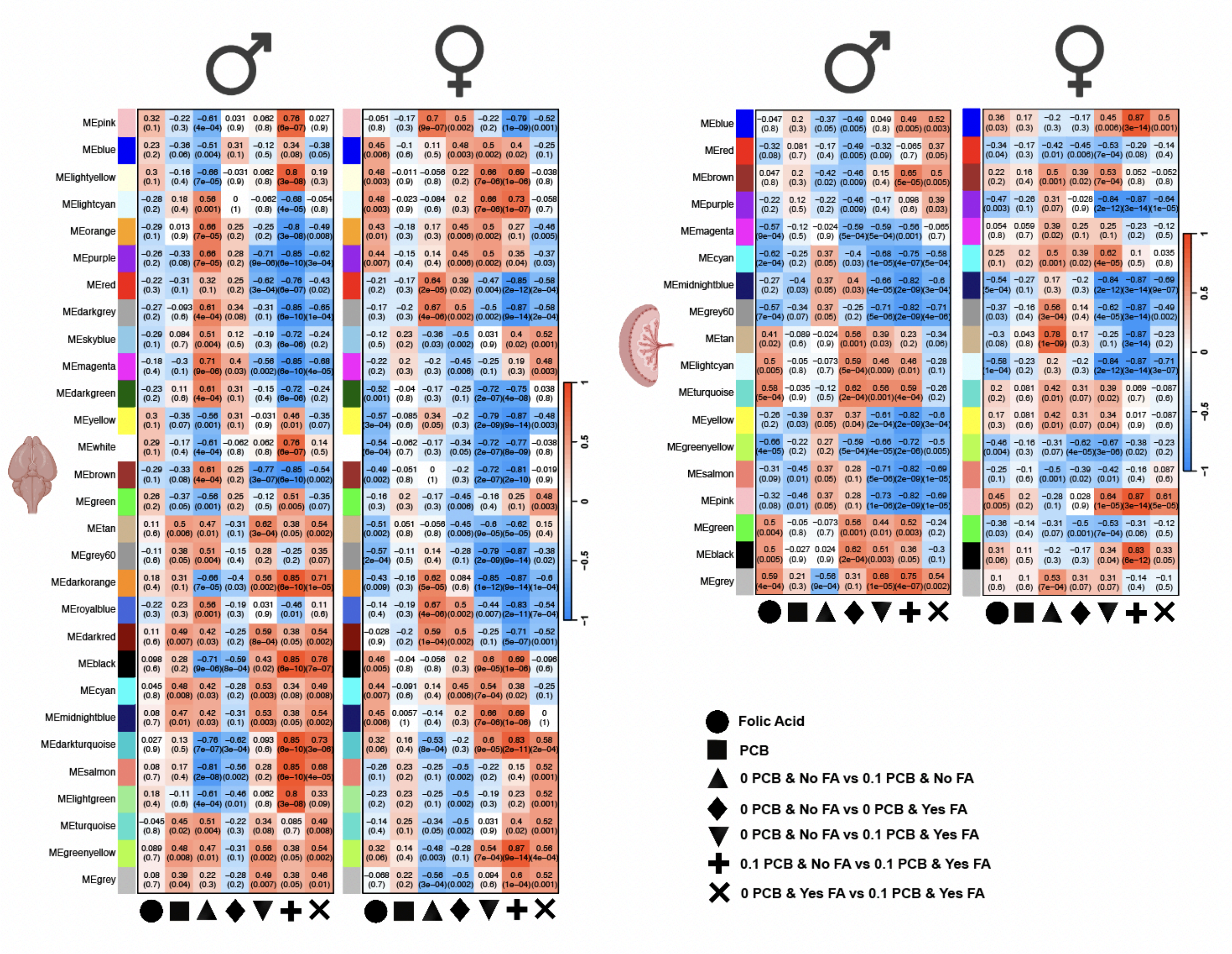


**Figure S9: Additional pairwise comparisons for sex-consensus folate networks.** This is an expanded version of Figure 6 but with all the pairwise comparisons.

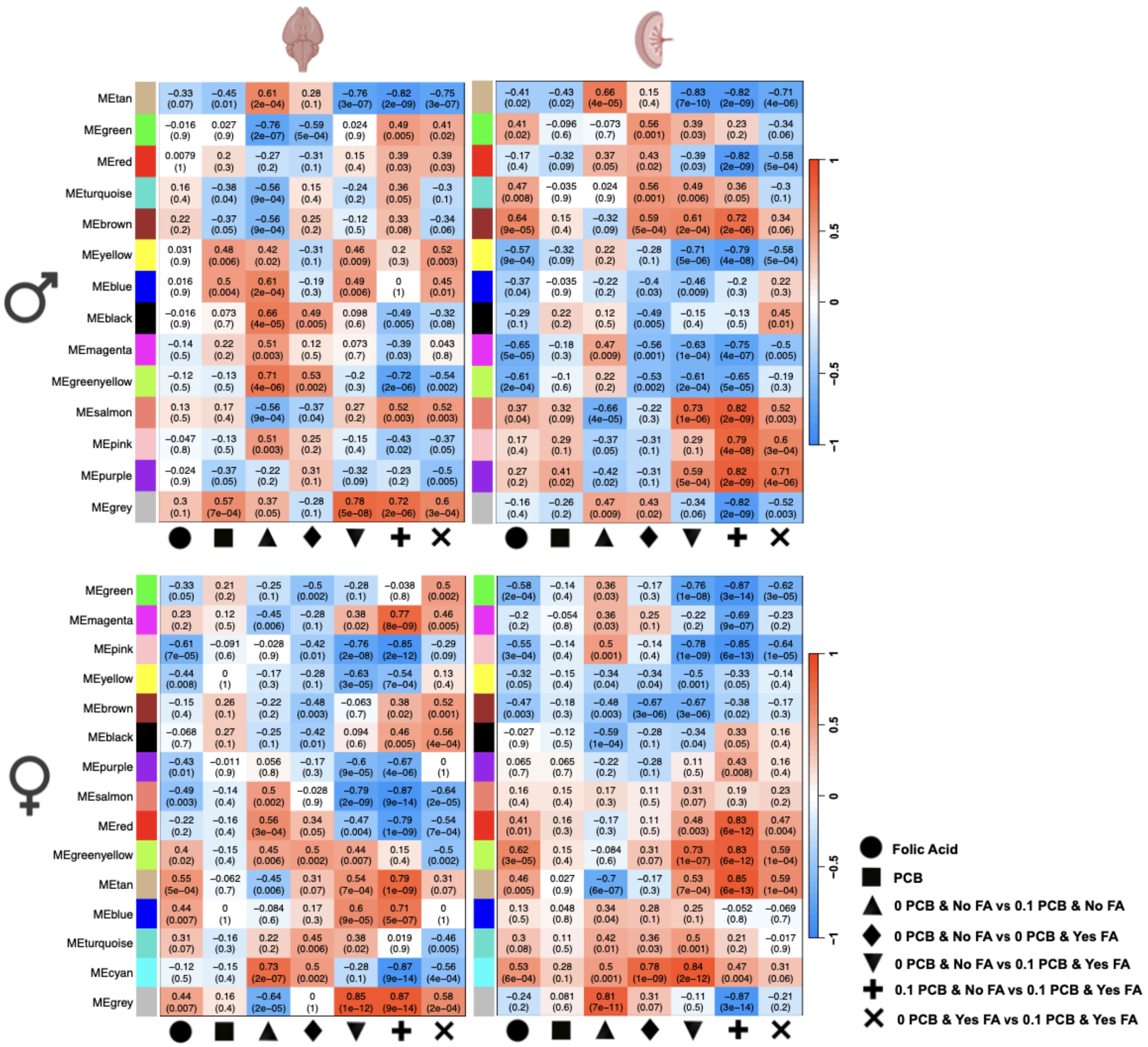


**Figure S10: Additional pairwise comparisons for tissue-consensus folate networks.** This is an expanded version of figure 7 but with all the pairwise comparisons.
